# Granule microenvironment regulates the dual functions of FMR1

**DOI:** 10.64898/2026.08.03.741171

**Authors:** Vaishali Grewal, Frank Wippich, Danilo Lüdke, Matteo Bordi, Annika Ladwig, Anna Orekhova, Anke Busch, Mandy Rettel, Julian König, Kathi Zarnack, Anne Ephrussi

## Abstract

Fragile X Messenger Ribonucleoprotein 1 (FMR1) is an evolutionarily conserved RNA binding protein with important functions in cognition and female reproduction, and its disruption is associated with neurodevelopmental and reproductive disorders including the Fragile X syndrome. FMR1 is best known for its role as a translation repressor. However, several recent studies also suggest a role of FMR1 as a translation enhancer raising fundamental questions about the molecular regulation of these opposing functions. In this study, we identify FMR1 as part of the *oskar* mRNA-protein complex in the *Drosophila* oocyte and study the role of FMR1 as a translational enhancer of *oskar*. We provide the molecular mechanism for the dual functions of FMR1 and show that the two major RNA-binding domains of FMR1, the KH domains and the RGG box, play distinct separable roles in regulating translation. The KH domains enhance translation of mRNAs while the RGG box containing C-terminal domain (CTD) is required to repress translation. We further show that the condensation propensity of FMR1 containing granules regulates the two antagonistic functions, such that phase separation by FMR1-CTD creates the molecular microenvironment necessary for the repressive activity, whereas reduction in phase separation is associated with increased translation. Our findings highlight the importance of biomolecular condensates not just as a means of molecular compartmentalization but as a fundamental regulatory principle that dictates the functional output of modular protein domains.

## Main

RNA binding proteins (RBPs) interact with and regulate various aspects of the RNA life cycle including transcription, localization, translation, and degradation^1^. Recent advances have highlighted the importance of the RNA - protein inter- and intra-molecular interactions in regulating the composition, physical state and in turn the functions of RNA - protein (RNP) complexes^2–4^. Altering the physical state of RNP granules leads to loss of function of constituent RNAs and proteins and is deleterious to growth and development^5–7^.

FMR1 is a versatile, highly conserved RNA binding protein with known roles in RNA transport, translation, granule formation and decay^8^. It comprises three RNA binding domains: two KH domains and an RGG box. Mutations in the *fmr1* gene are known to cause severe disorders in humans, such as Fragile X primary ovarian insufficiency (FXPOI), Fragile X-associated tremor and ataxia syndrome (FXTAS) or Fragile X-associated neuropsychiatric disorder (FXAND)^9^. In extreme cases, mutations in *fmr1* cause Fragile X Syndrome (FXS), an inherited form of mental disability^10^.

Due to the important, conserved roles of FMR1 in RNA regulation, several studies have focused on understanding the molecular basis of its various functions. The protein has been predominantly identified as a translation repressor of its target RNAs^11–17^. FMR1 directly associates with ribosomes using its KH domains^12^, an interaction previously thought to be crucial for its repressive activity. Recent studies, however, have indicated that the C-terminal RGG box is necessary and sufficient for repression^18, 19^, and that the phase separation propensity of the RGG box is crucial for the repressive activity of FMR1^17, 20^. Besides translation repression, there is also accumulating evidence to support a role of FMR1 and its paralogs FXR1 and FXR2 in translation stimulation^21–28^. However, the molecular mechanism of translation stimulation by FMR1, and how the two antagonistic functions of the protein are regulated, remains unclear.

In this study, we show that FMR1 is a novel component of *oskar* mRNA-containing RNP granules (mRNPs) in *Drosophila* oocytes and acts in positively regulating Oskar protein levels. We dissect the molecular mechanism underlying the opposing translation enhancing and repressing functions of FMR1 and show that the KH domains enhance translation while the C-terminal RGG box inhibits translation. Additionally, FXS causing mutations in KH domains (I244N and I307N) that impair RNA and ribosome binding abolish translation stimulation, indicating a role of these residues in the translation mechanism. We further demonstrate that the condensation propensity of FMR1 containing granules acts as a molecular switch between its two functions such that phase separation of the FMR1 C-terminal domain is associated with translation repression, whereas loss of condensation favors translation.

## Results

### Transcript-specific pulldown identifies FMR1 as a novel component of *oskar* mRNP granules

Proper embryonic patterning in *Drosophila melanogaster* depends on the correct spatial and temporal expression of maternal mRNAs in the *Drosophila* oocyte^29–32^. *oskar* mRNA is one such essential maternal mRNA, transcribed in the nurse cell nuclei and actively transported to the posterior of the developing oocyte. Once localized, the mRNA is translated into Oskar protein, which is essential for embryonic patterning and for the initiation of pole plasm formation^31, 33, 34^. *oskar* mRNA associates with several proteins, forming mRNP granules which tightly regulate its localization and translation, making it an ideal model for the study of RBP function and regulation in a native, granule context. Though the identities and functions of many RBPs involved in *oskar* regulation have been discovered over the years, a comprehensive atlas of the proteins that associate with *oskar* granules is still lacking. Therefore, to identify the RBPs associated with *oskar* in an unbiased manner, we performed an *oskar*-specific pulldown, as described in Wippich and Ephrussi 2020^35^. Briefly, mRNP complexes were crosslinked *in vivo* using UV irradiation and formaldehyde, followed by isolating the complexes using biotinylated probes complementary to the target RNA - here *oskar* RNA. The enriched proteins were then analyzed by mass spectrometry (Figure 1a). We observed enrichment of proteins previously identified to be a part of *oskar* mRNP granules, including Bruno (Bru/ARET), Me31B, Stau, Hrp48 and PTB (Supplementary Figure S1). Additionally, we enriched for proteins with no previously known role in *oskar* regulation, among which was the translational regulator FMR1 (Figure 1b). Considering the important roles of FMR1 in RNA regulation and human physiology, we decided to further investigate its role as an *oskar* granule component in the germline.

**Figure 1:**
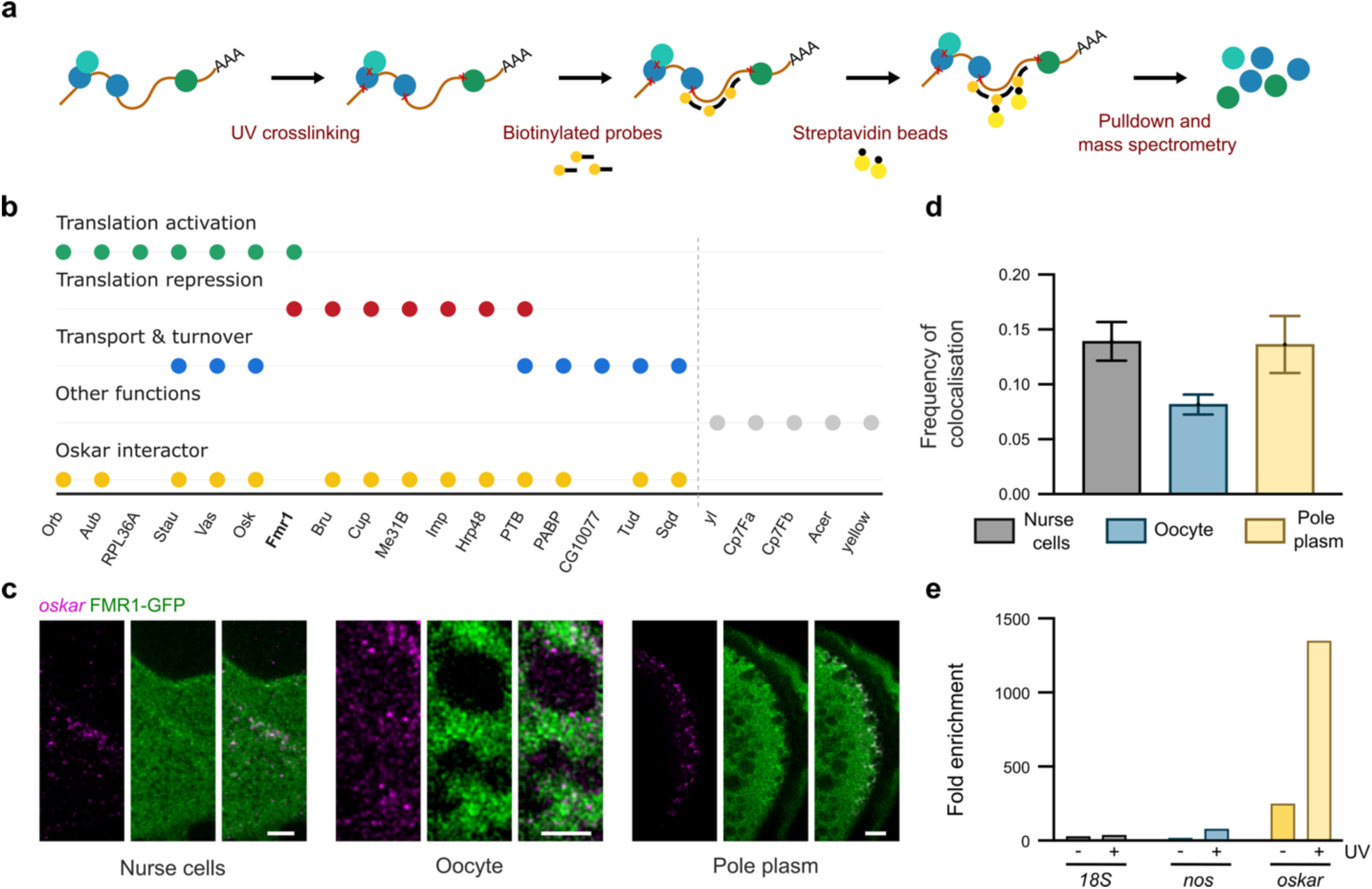
Transcript-specific pulldown of *oskar* identifies FMR1 as a novel component of the *oskar* mRNP granules. **a**. Schematic of *oskar*-specific pulldown to identify the protein components of *oskar* granules. **b**. Classification of enriched protein based on functions and previously known interactions with *oskar*. **c**. Co-localization analysis between FMR1-GFP and *oskar* mRNA in the nurse cells, the oocyte and the pole plasm (scale bar – 5 µm). **d**. Quantification of the frequency of co-localization between FMR1-GFP and *oskar* in the different compartments. Error bars represent standard error. **e**. CLIP experiment showing fold enrichment of *oskar* mRNA with FMR1-GFP upon UV crosslinking. *18S* and *nos* RNAs were used as controls.

### FMR1 and *oskar* mRNA interact directly *in vivo*

To validate the putative interaction of *oskar* mRNA with FMR1 identified by our transcript-specific pulldown, we performed an object-based co-localization analysis. *oskar* mRNA was labeled by single molecule fluorescent *in situ* hybridization (smFISH) in egg chambers of flies expressing FMR1-GFP. We observed that FMR1-GFP showed a significant co-localization with *oskar* mRNA in the nurse cells, within the oocyte, as well as at the oocyte posterior pole (Figure 1c and 1d). This indicates that FMR1 is indeed associated with *oskar* granules *in vivo* in all compartments, from the nurse cells to the oocyte posterior pole.

Furthermore, since the transcript-specific pulldown relies on both UV and formaldehyde crosslinking, which capture both direct and indirect protein - RNA interactions and protein - protein interactions, we sought to determine whether the enrichment of FMR1 with *oskar* mRNA was a result of direct binding of FMR1 to *oskar,* or of indirect binding, via protein - protein interactions. To test this, we performed a UV crosslinking and immunoprecipitation (CLIP) experiment in flies expressing FMR1-GFP. The RNA-FMR1-GFP complexes were UV crosslinked *in vivo* and pulled down using an anti-GFP nanobody. The enriched RNAs were analyzed by qRT-PCR for the presence of *oskar* mRNA. As compared to a non-crosslinked sample, *oskar* mRNA was significantly enriched with FMR1-GFP in the UV crosslinked samples (Figure 1e). This indicates that FMR1-GFP directly binds to *oskar* mRNA *in vivo*.

### iCLIP identifies specific FMR1 binding sites on *oskar* mRNA

With evidence for a direct interaction of *oskar* with FMR1 *in vivo*, we sought to determine the binding sites of FMR1 across the transcriptome at a nucleotide resolution. To this end, we performed iCLIP (individual-nucleotide resolution CLIP)^36^ on ovaries from flies expressing FMR1- GFP (Figure 2a, left). iCLIP relies on UV-induced covalent crosslinking of RNA binding proteins to their target RNAs, followed by immunoprecipitation of the complexes. During reverse transcription, truncation at the crosslinked nucleotide enables mapping of protein - RNA interactions at single-nucleotide resolution. We performed iCLIP on three sample types - FMR1- GFP, GFP control, and beads-only control - with four replicates each.

**Figure 2:**
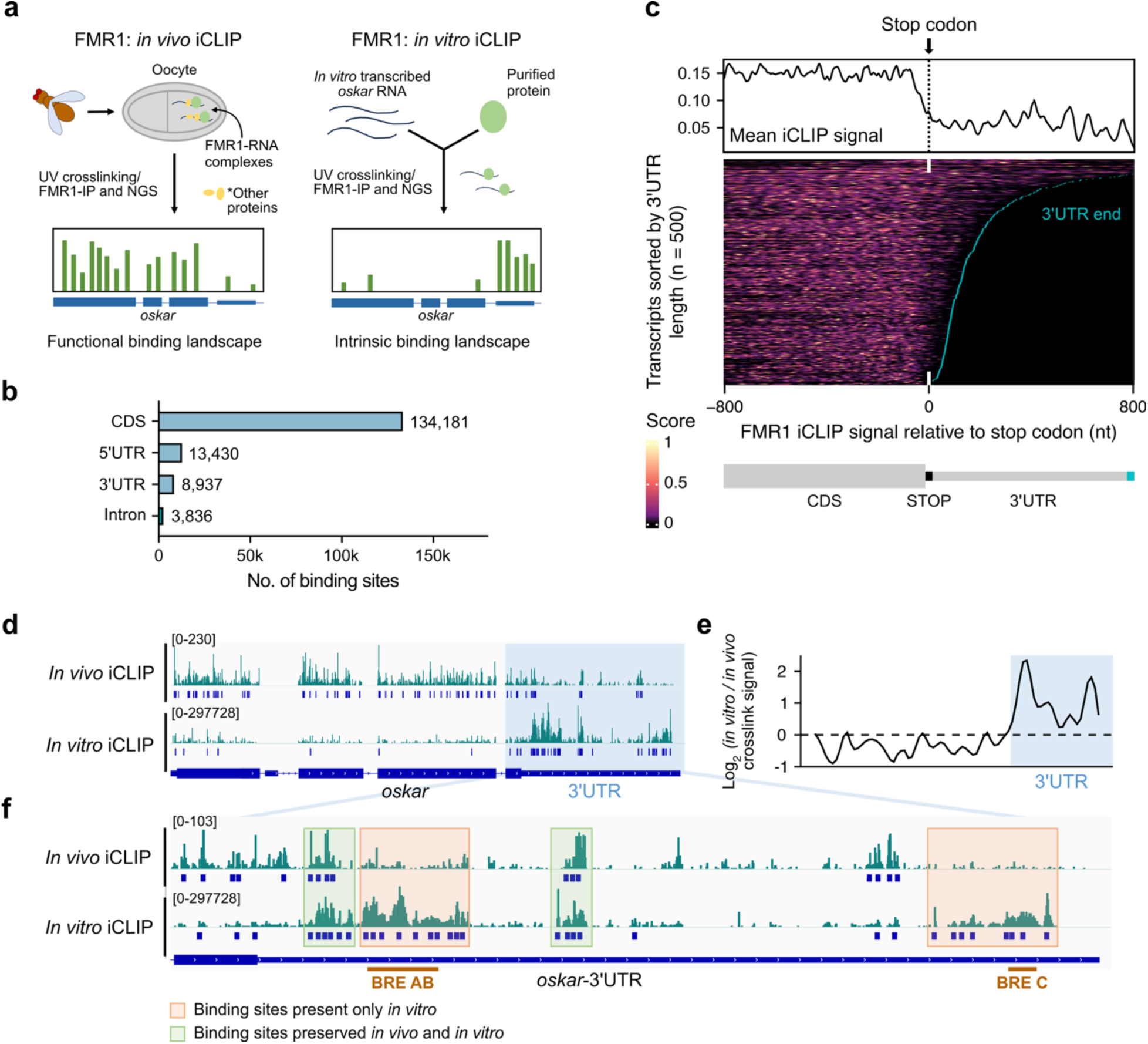
iCLIP identifies binding sites of FMR1 on *oskar* mRNA. **a**. Schematic of *in vivo* and *in vitro* iCLIP of FMR1 in the experimental setup. **b**. Graph shows the distribution of FMR1 binding sites across the transcript regions. **c**. Heatmap of normalized (row-wise min-max normalization) crosslink signal *in vivo* centered on transcript stop codons. The 500 transcripts with highest cumulative crosslink signal within the displayed region were aligned at the stop codon (0 nt) and sorted by decreasing 3’UTR length. Color intensity ranges from black (0; low/no signal) to yellow (1; high signal). Cyan lines denote annotated 3’UTR ends. The meta profile above the heatmap represents the mean normalized crosslink signal across all displayed transcripts, excluding positions outside annotated transcript boundaries. (Note: normalized crosslink signal score was squared) **d**. FMR1 crosslink signal and identified binding sites across the full-length *oskar* transcript *in vivo* (top panel) and *in vitro* (bottom panel). **e**. Smoothed log2 ratio of Z-score-normalized crosslink signal along *oskar* mRNA, comparing *in vitro* to *in vivo* signal. Positive values indicate relatively higher *in vitro* signal, whereas negative values indicate relatively higher *in vivo* signal. The annotated 3’UTR is highlighted in light blue. **f**. FMR1 crosslink signal and identified binding sites across the *oskar* 3’UTR *in vivo* (top panel) and *in vitro* (bottom panel). The two binding sites for Bruno (BRE AB and BRE C) are marked. Highlighted regions indicate shared (light green) and *in vitro*-enriched (light orange) binding sites. For *in vivo* iCLIP n=4, for *in vitro* iCLIP n=3 for 250 nM ΔN-FMR1-GFP and n=4 for 100 nM ΔN-FMR1-GFP.

To identify crosslink signal above background and determine significant binding sites, we used PureCLIP for peak calling^37^. Binding sites were defined as clusters of significant crosslink sites spanning seven nucleotides using BindingSiteFinder^38^. We recovered 160,960 reproducible binding sites across the transcriptome present in at least three out of four replicates (Figure 2b and S2a). The binding sites occurred in the transcripts of 3,677 genes, the majority of which were protein-coding (Figure S2b).

Within the bound transcripts, most binding sites (∼83%) were located in the coding region, with only 14% in the 5′ and 3′ untranslated regions (UTRs; Figure 2b). A similar enrichment of FMR1 within coding regions has been observed in CLIP and TRIBE studies in mouse and *Drosophila*^13,39^. Notably, in meta profiles, FMR1 binding was enriched across the entire open reading frame of its target transcripts, with a pronounced decline in crosslink signal immediately downstream of the stop codons, marking a clear transition from the coding sequence (CDS) into the 3′UTR (Figure 2c). This pattern suggests that FMR1 *in vivo* is closely associated with actively translated regions of transcripts, consistent with previous findings linking FMR1 to translating ribosomes^12–14^. In line with the *oskar*-transcript pull-down assays, *oskar* mRNA was identified as a prominent target of FMR1 in the iCLIP data (Figure 2d). Consistent with the transcriptome-wide binding pattern, significant FMR1 binding sites were distributed throughout the coding region. In addition, several pronounced binding sites were detected in the 3′UTR (Figure 2e).

To distinguish binding events driven by the intrinsic RNA-binding affinity of FMR1 from those shaped by cellular factors, we next performed *in vitro* iCLIP^40^ to determine the inherent binding of FMR1 on *oskar* mRNA (Figure 2a, right). In contrast to *in vivo* iCLIP, which captures the physiological binding landscape in the cellular context, *in vitro* iCLIP reveals intrinsic RNA-binding preferences independent of additional cellular components. Experiments were performed using recombinantly purified FMR1 (ΔN-FMR1-GFP, amino acids 220–681, containing all RNA-binding domains) and full-length *in vitro* transcribed *oskar* mRNA (Figure 2a).

Direct comparison of *in vivo* and *in vitro* crosslink profiles across the *oskar* mRNA revealed distinct binding behaviors within the CDS and the 3’UTR (Figure 2d, e). In the CDS, FMR1 binding differed markedly between the two conditions. Whereas strong FMR1 binding was observed throughout the coding sequence of *oskar* mRNA *in vivo*, this was largely lost under *in vitro* conditions. In contrast, the binding pattern within the 3′UTR was more differentiated. While several binding sites were preserved between *in vivo* and *in vitro*, indicating intrinsic recognition by FMR1, others showed enhanced or reduced binding *in vitro*, reflecting the absence of factors that modulate FMR1 binding *in vivo* (Figure 2f). Notably, a subset of sites was specifically bound *in vitro*. These correspond to the Bruno response element (BRE) AB and BRE C sites - regions normally occupied by Bruno, a key translational repressor of *oskar* during its transport from nurse cells to the oocyte. The emergence of these binding sites *in vitro* therefore points to competitive binding of Bruno and FMR1 in the *oskar* 3′UTR.

Together, these findings indicate that FMR1 can directly bind to specific sites within the *oskar* 3′UTR, and that its binding is subject to regulation by additional *trans*-acting factors *in vivo*. In addition, the strong difference in the CDS suggests that the extensive association of FMR1 with coding regions is not primarily driven by its intrinsic RNA-binding specificity, supporting the proposed association of FMR1 with translating ribosomes.

### Loss of FMR1 leads to reduced Oskar protein levels *in vivo*

Since FMR1 is a known translational regulator, we wanted to study if the loss of FMR1 has any effect on Oskar protein levels. To this end, we analyzed Oskar protein levels in *fmr1* loss of function mutants (Fmr1^Δ50M^/Fmr1^Δ113M^), as well as in flies in which FMR1 was knocked down specifically in the germline. The knockdown was performed using the germline specific *oskar*- Gal4 to drive the expression of shRNA in FMR1-RNAi line. In both cases, we observed a significant, 50% reduction in the levels of both the long and the short Oskar isoforms (Figure 3a and 3b). To see if this might be due to a reduction in *oskar* RNA levels, we performed a qRT-PCR for *oskar* in FMR1-RNAi versus control (*white)* RNAi flies and observed no significant difference (Supplementary Figure S3a). This indicates that FMR1 positively regulates Oskar protein levels in the oocyte, consistent with the less studied role of FMR1 in enhancing translation.

**Figure 3:**
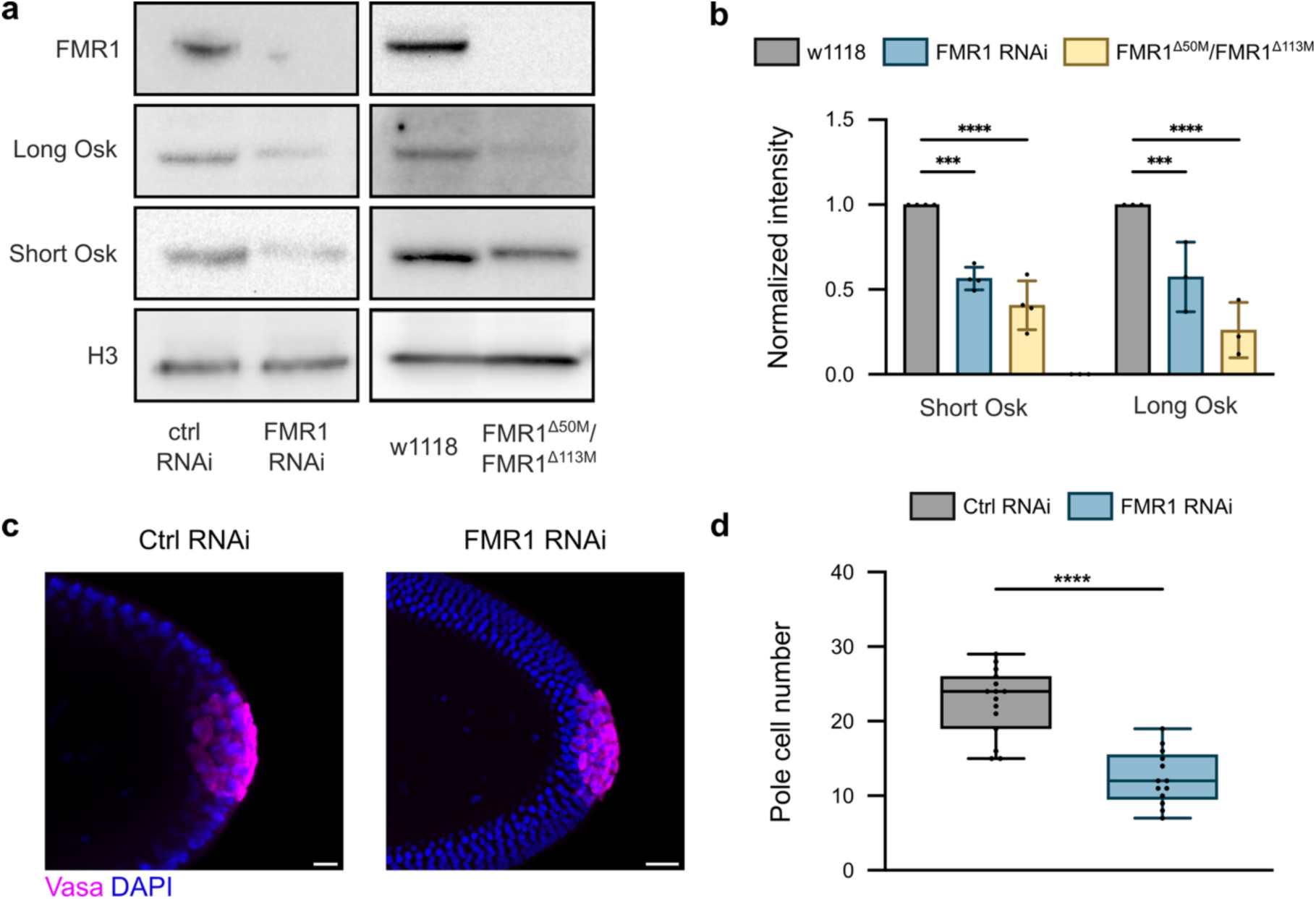
Loss of FMR1 leads to reduced Oskar protein levels *in vivo*. **a**. Western blot to detect Oskar protein levels in FMR1 knockdown and loss of function mutants (FMR1^Δ50M^/FMR1^Δ113M^) (Note: FMR1 staining (top panel) was performed in an independent experiment to confirm functional RNAi line and mutants) **b**. Graph shows the quantification of decrease in Oskar protein. One-way ANOVA test was used to determine the statistical significance. Each dot represents a replicate. Error bars represent standard deviation. P-values are ****<0.0001, ***<0.001. **c**. Pole cells (stained using Vas antibody) in control (*white*) RNAi vs FMR1 RNAi (scale bar – 10 µm). **d**. The graph shows the number of pole cells in each case. Each dot represents one embryo. Student’s *t*-test was used for statistical analysis, and P-value is <0.0001.

Oskar protein is essential for the induction of pole cell formation and abdominal patterning in a dose-dependent manner, such that partial knockdown of *oskar* affects pole cell formation, without affecting abdominal patterning^41^. Consistent with the reduced amount of Oskar in the ovaries of FMR1 knockdown flies, we observed that the number of pole cells produced by the FMR1 knockdown embryos was approximately 50% of that produced by the control (*white*) RNAi flies (Figure 3c and 3d). This is in agreement with the finding of Deshpande et al. 2006^42^, who also observed a reduced number of pole cells in embryos produced by *fmr1*^3^/ *fmr1*^3^ mutant flies. The reduction in Oskar protein, however, was not sufficient to affect hatching rate or abdominal patterning, as expected from the dose-dependent role of *oskar* (Supplementary Figure S3b and S3c).

### KH domains of FMR1 stimulate translation of reporter mRNA *in vitro*

To systematically study the effect of the two RNA binding domains of FMR1 on translation of RNAs, we performed an *in vitro* translation assay using *Drosophila* embryo lysates. We used the bacteriophage **λ**N-BoxB interaction^43, 44^ to tether **λ**N-sfGFP tagged FMR1 domains to luciferase reporter RNAs containing BoxB elements in the 3’UTR (luc_BoxB) and analyzed the effect on the translation of reporters (Figure 4a). We used two protein constructs at 8 µM concentration: λN- sfGFP-FMR1-KH that contains only the two KH domains, and λN-sfGFP-FMR1-CTD that contains only the RGG box-containing C-terminal domain (CTD) of FMR1.

**Figure 4:**
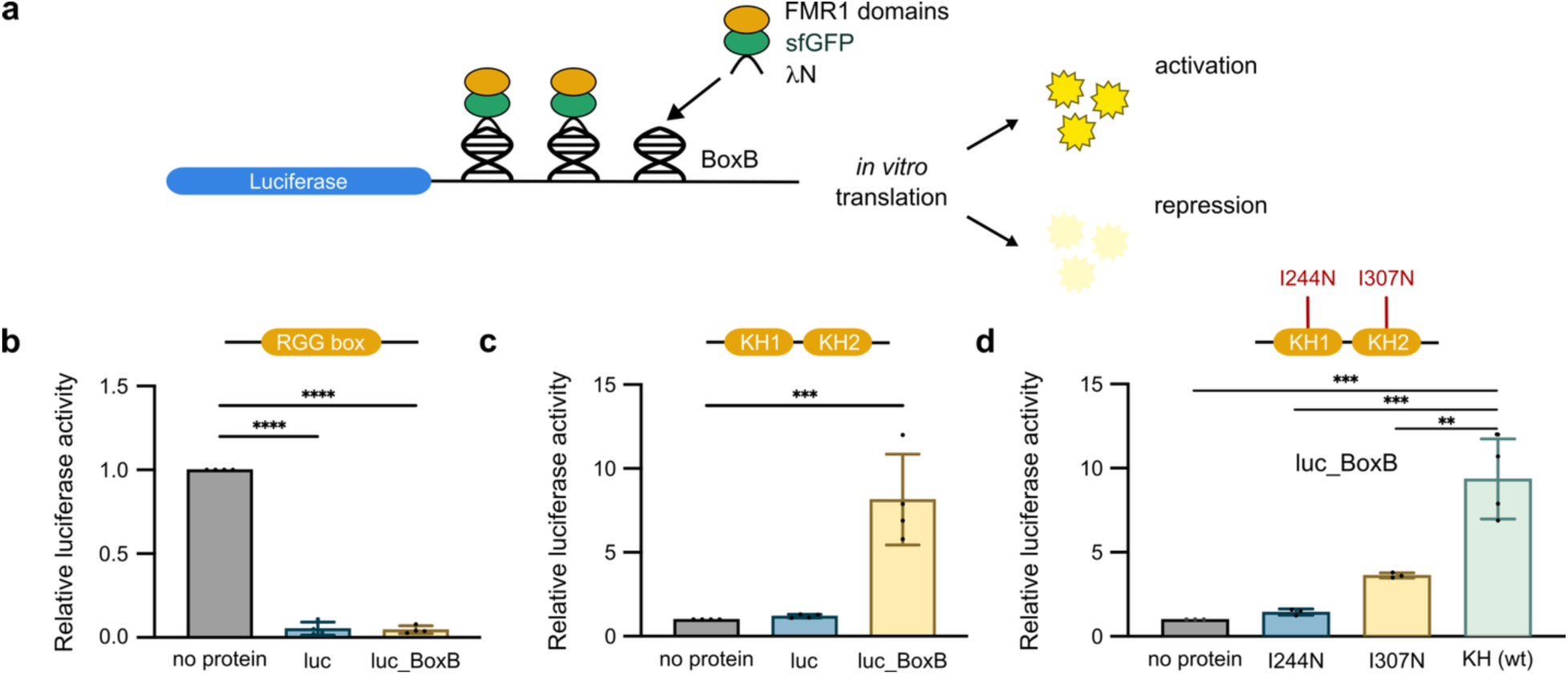
Domain dependent effect of FMR1 on translation of reporters. **a**. Schematic of reporter and proteins used for the *in vitro* translation assay. **b**. and **c**. show graphs with the relative luciferase activity upon FMR1 tethering (domains tethered are labeled on top in orange) as compared to no protein control. **d**. Effect of KH domain mutants I244N and I307N on translation of luc_BoxB reporter. KH(wt) is the wildtype domain from panel **c**. Error bars represent standard deviation. One-way ANOVA test was used for statistical analysis. Each dot represents a biological replicate. P-values: ****<0.0001, ***<0.0005, **<0.005.

We observed that when FMR1-CTD is added to the reporters, translation of both luc and luc_BoxB reporters was strongly repressed (Figure 4b). This is consistent with recent studies showing that the FMR1-CTD is sufficient for translation repression^17–20^. FMR1-KH, however, had the opposite effect and caused a significant increase in translation of the reporter luc_BoxB as compared to control luc reporter (Figure 4c). FMR1 thus seems to have a domain dependent effect on the translation of reporter RNAs upon tethering, wherein the C-terminal domain represses translation, and the KH domains stimulate translation. We believe that CTD represses the control luc as well as the luc_BoxB because RGG box has degenerate specificity for RNA binding^45^ as compared to KH domains which bind more specifically^46^. FMR1-CTD is therefore able to interact with the reporter RNA without requiring BoxB and represses translation at 8 µM.

### FXS associated mutations in KH domains abolish translation stimulation

Mutations in the KH domains of FMR1 are associated with impaired protein function and disease-associated phenotypes in humans. In particular, I241N in KH1 is known to disrupt actin dynamics^47^ and axonal development^48^, and I304N in KH2 is associated with the Fragile X Syndrome (FXS)^49^. These residues are highly conserved^50^ and the corresponding mutations in *Drosophila* FMR1, I244N and I307N, cause abnormalities in mushroom body axon guidance, disrupted courtship behavior, and altered circadian rhythm^51^. At the molecular level, both mutants show disrupted RNA binding and granule formation^52^, and reduced association with ribosomes and polysomes^12, 14, 19^. To test the effect of the KH domain mutants on translation, we cloned and purified the FMR1-KH(I244N) and FMR1-KH(I307N) proteins and assessed their function in the *in vitro* translation assay.

We observed that KH(I244N) completely failed to stimulate translation of the luc_BoxB reporter, and KH(I307N) showed significantly reduced translation when compared to wildtype KH domains (Figure 4d). These findings indicate that mutations in key KH domain residues impair the translation stimulation function of the protein. Our results also suggest that disruption of KH domain-mediated translational stimulation may contribute to the molecular dysfunction underlying FXS and other FMR1-associated disorders, requiring further investigation.

### KH domains are sufficient to maintain normal Oskar levels *in vivo*

As the KH domains are necessary and sufficient for translation *in vitro*, we hypothesized that FMR1 containing only the KH domains should be sufficient to maintain normal Oskar protein levels *in vivo*. To test this, we used CRISPR-Cas9 mediated mutagenesis to generate a fly line in which the endogenous C-terminal domain of FMR1 was deleted and replaced by an sfGFP tag (hereafter called FMR1-KH-GFP). Additionally, we tagged the endogenous full length FMR1 with sfGFP as control (hereafter called FMR1-sfGFP). Expression of the sfGFP tagged proteins was verified by microscopy as well as western blot analysis (Figure 5a and 5b). We observed Oskar protein levels in FMR1-KH-GFP are comparable to FMR1-GFP and w1118 (control) (Figure 5b), Consistent with the normal Oskar levels, the number of pole cells in FMR1-KH-sfGFP and FMR1- sfGFP expressing fly lines were also comparable to w1118 (control) (Figure 5c and 5d). This confirms that the KH domains of FMR1 are indeed necessary and sufficient to maintain Oskar protein levels *in vivo*.

**Figure 5:**
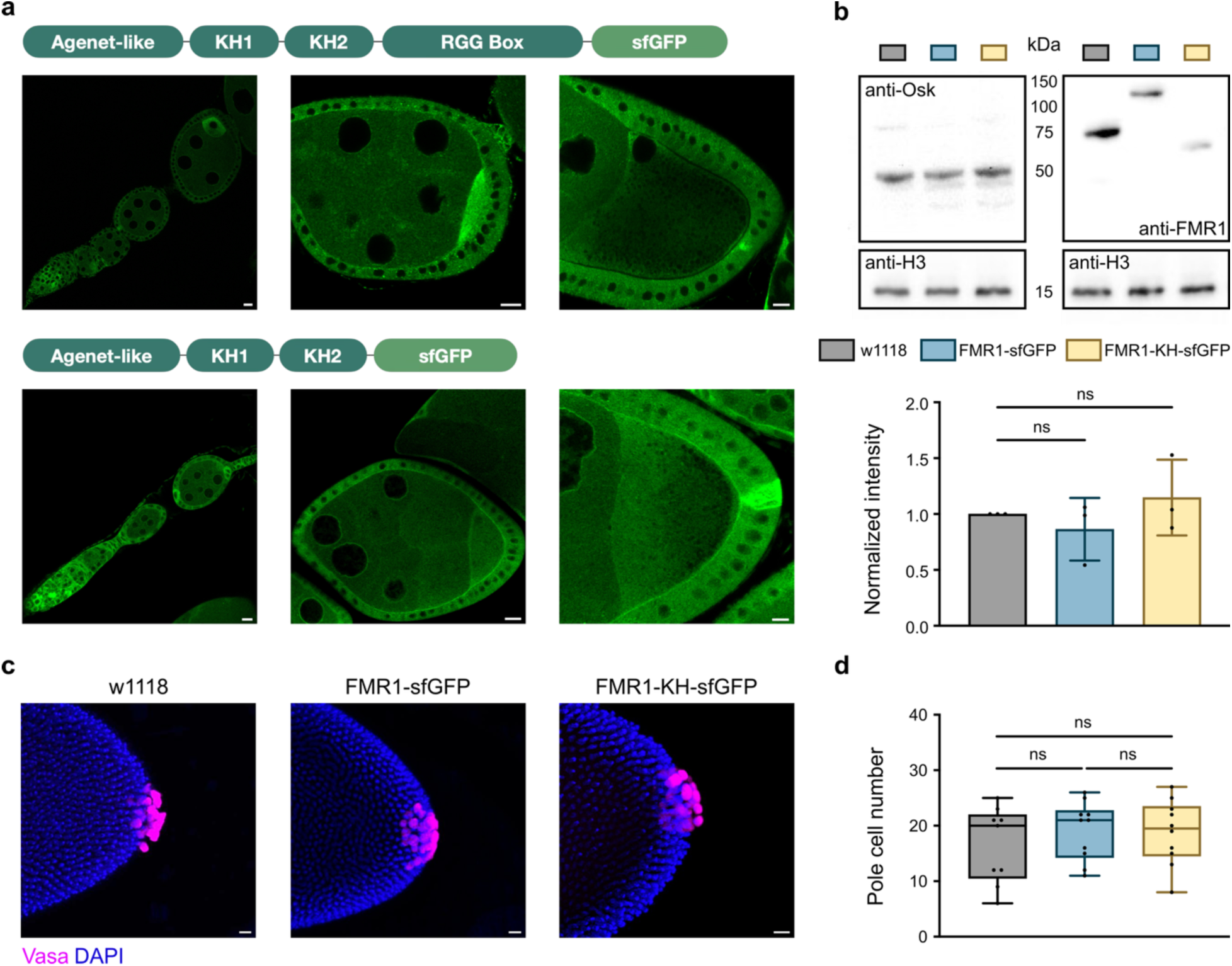
KH domains are sufficient to maintain normal Oskar levels *in vivo*. **a**. Expression of FMR1-sfGFP and FMR1-KH-sfGFP in fly lines generated by CRISPR Cas9 mutagenesis. **b**. Western blot to detect Oskar protein levels in w1118, and CRISPR generated FMR1-sfGFP and FMR1-KH-sfGFP. Graph shows the quantification of band intensity relative to H3. Error bars represent standard deviation. Each dot represents a biological replicate. One-way ANOVA test was used for statistical analysis. **c**. Pole cells (stained using Vas antibody) in w1118, FMR1-sfGFP and FMR1-KH-sfGFP lines (scale bar – 10 µm). **d**. Graph shows the number of pole cells in each case. Error bars represent standard deviation. Each dot represents one embryo. One-way ANOVA test was used for statistical analysis.

### The physical state of granules regulates the dual activity of FMR1

We next sought to understand how the two opposing functions of FMR1 are regulated. Studies have shown that the phase separation propensity of the FMR1-CTD has an important role in translation repression^17, 20^. Consistent with this, visualizing the *in vitro* translation mixtures for λN- sfGFP-FMR1-KH and λN-sfGFP-FMR1-CTD under microscope revealed phase separated granules with only λN-sfGFP-FMR1-CTD, evident in both DIC (differential interference contrast) and fluorescent images (Figure 6a). Analysis of the size and circularity of the granules showed that FMR1-CTD formed significantly larger granules with higher circularity as compared to FMR1- KH (Figure 6a).

**Figure 6:**
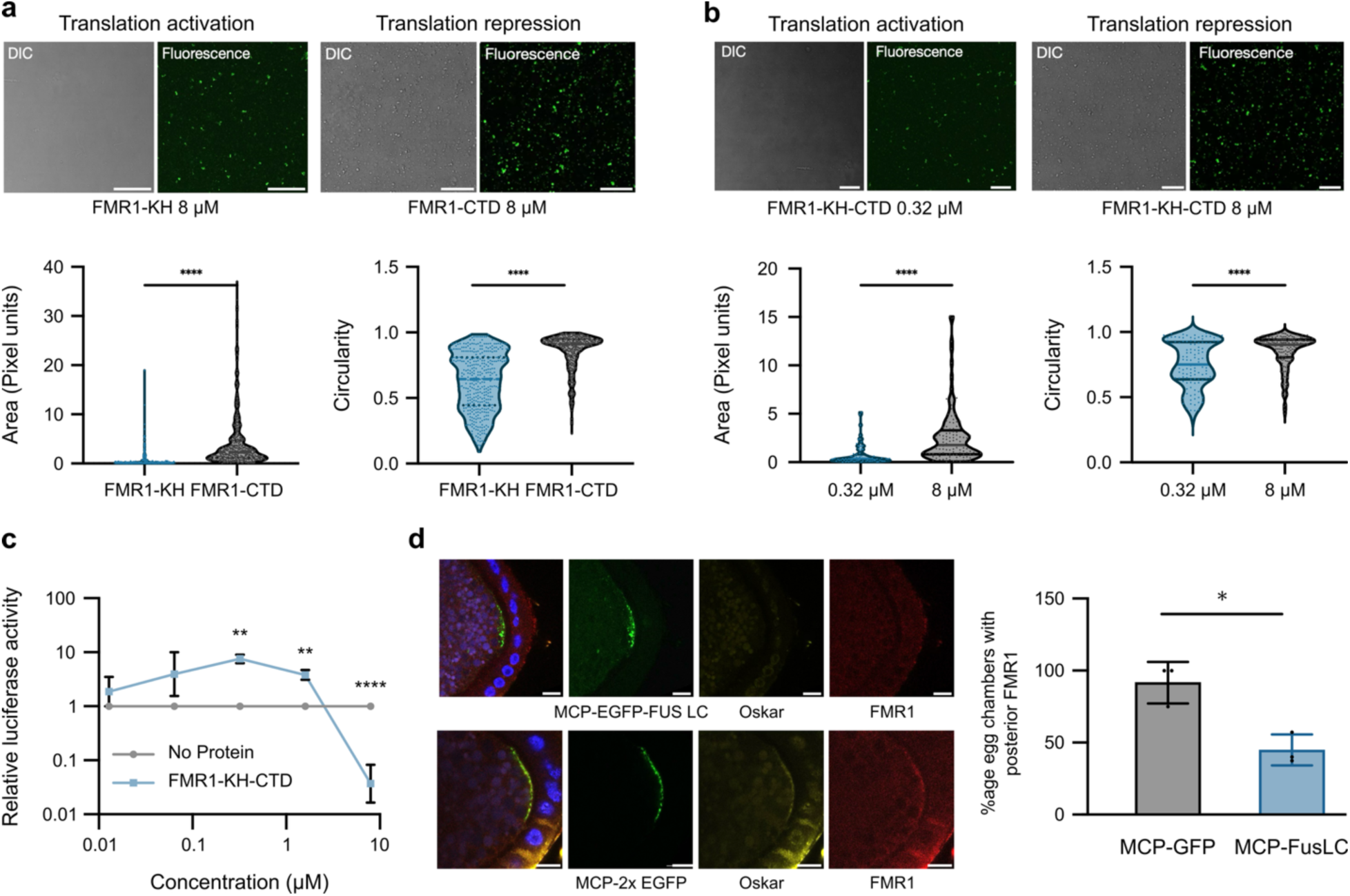
The physical state of granules regulates the dual activity of FMR1. **a**. *In vitro* translation reaction mixtures with 8 µM λN-sfGFP-FMR1-KH and λN-sfGFP-FMR1-CTD observed under microscope with differential interference contrast (DIC) image of granules on the left and fluorescence image on the right. Graphs show the size and circularity of the granules formed by the two proteins. Each dot represents one granule. **b**. *In vitro* translation reaction mixtures with 0.32 µM and 8 µM λN-sfGFP-FMR1-KH-CTD. Graphs show the size and circularity of the granules formed by the two proteins. **c**. Graph shows the effect of different concentrations of λN-sfGFP-FMR1-KH-CTD on translation of luc_BoxB reporter normalized to no protein control. **d**. Immunostaining of Oskar and FMR1 in flies expressing MCP-EGFP-FUS LC and MCP-2X EGFP (control) (scale bar - 10 µm). Graph represents the percentage of egg chambers showing enrichment of FMR1 at the posterior pole in MCP-EGFP-FusLC and MCP-2xEGFP (control) conditions. Error bars represent standard deviation. Each dot represents one biological replicate. 5-15 egg chambers counted for each replicate. Student’s *t*-test was used for statistical analysis. P-values: ****<0.0001, ***<0.0005, **<0.005.

To investigate the effect of the FMR1 on translation when both domains are present, we performed the *in vitro* translation assay with λN-sfGFP-FMR1-KH-CTD. We performed the experiment across a series of five-fold dilutions, while simultaneously monitoring condensate formation. We found that, like FMR1-CTD, FMR1-KH-CTD exhibits evident phase separation and represses translation at 8 µM. However, subsequent dilutions led to an increase in the translation of the reporter, with a maximum increase in fold change at 0.32 µM, followed by no significant stimulation/repression (Figure 6c). Additionally, no phase separation was observed at 0.32 µM, and the granules were significantly smaller and had lower circularity as compared to the 8 µM condition (Figure 6b). This indicates that at low concentrations insufficient to induce phase separation, tethering FMR1 to target RNAs can stimulate translation, but as the concentration increases and phase separation, is triggered the activity switches to a repressive function. It is therefore plausible that the translation output of FMR1 depends on the types of proteins and granules with which FMR1 associates. In contexts where translation is required, for instance upon localization of *oskar* mRNA, FMR1 must interact with proteins and granules within a non-phase separated, translation permissive microenvironment (Figure 6a, 6b and 6c).

Our previous work has shown that *oskar* forms solid granules *in vivo*, and “liquification” of these granules, by tethering the Fus low complexity domain (FUS LC) to *oskar* via the MCP-MS2 interaction, leads to reduced translation of *oskar* mRNA^7^. To assess if altered association and/or activity of FMR1 contributes to the loss of *oskar* translation upon FUS LC tethering, we immunostained for FMR1 and analyzed its association with the phase altered *oskar-*FUS granules. We observed that FMR1 fails to localize with the *oskar*-FUS granules and loses the posterior enrichment observed under control (*oskar*-MCP-EGFP) conditions (Figure 6d). Altering the state of *oskar* granules therefore alters their RNP composition and leads to failure of FMR1 association, suggesting that disrupted FMR1 recruitment contributes to the loss of *oskar* translation.

## Discussion

In this study we identified FMR1 as a novel protein component of *oskar* mRNP granules and found that, contrary to its traditional role in repressing translation, FMR1 has a positive effect on Oskar protein levels. This led us to uncover a domain dependent mechanism of how FMR1 performs its dual, antagonistic roles in regulating translation. We found that the KH domains of FMR1 have a role in stimulating translation and are dispensable for its repressive activity. In contrast, the CTD has a strong repressive effect on translation. Furthermore, disease causing I244N and I307N mutations in KH domains are crucial for its role in translation stimulation.

Our findings suggest that the dual activities of FMR1 are regulated not only by the two RNA binding domains, but also by the granule microenvironment. Given that CTD induced liquid-liquid phase separation (LLPS) modulates the repressive activity of FMR1^17, 20^, the presence of interactors and post-translational modifications that either stimulate or repress LLPS of FMR1 containing granules would be crucial to switch between the two activities. In order to stimulate the translation of RNA targets such as *oskar*, FMR1 would therefore need to be maintained in a non-phase separated solid state, which is the case for *oskar* granules^7^. Tethering of FUS to *oskar* changes the microenvironment of the granules altering the granule composition and state. This affects the function of constituent proteins, including FMR1, which is unable to bind and stimulate translation of *oskar*. This suggests that FMR1 and its interactors *in vivo* fine tune the association of FMR1 within granules in different cellular contexts (for e.g., transport granules, stress granules, P-bodies)^8, 53, 54^ and determine the functions it will henceforth perform.

Our analysis of *oskar* and FMR1-GFP co-localization shows that FMR1 is a component of *oskar* mRNP granules in the nurse cells. The fact that FMR1 does not stimulate translation of *oskar* within the solid granules during transport suggests that other, dominant factors keep *oskar* in a translationally repressed state. A key *oskar* repressor is the protein Bruno, which is a core component of *oskar* granules and is also present in the nurse cells. Comparative analysis of our *in vivo* and *in vitro* iCLIP data reveals differences in FMR1 binding to certain sites in the *oskar* 3’UTR. For instance, in the *in vitro* iCLIP, FMR1 shows enhanced binding to Bruno binding sites BRE AB (Figure 2F, lower panel). Our *in vitro* competition experiments (Supplementary Figure S2c) show that Bruno, when added in increasing concentrations, outcompetes FMR1 for binding to BRE AB (Supplementary Figure S2d), whereas FMR1 is not able to outcompete Bruno (Supplementary Figure S2e), indicating that Bruno has a higher affinity than FMR1 for those sites. We speculate that Bruno binding keeps the translation of *oskar* repressed during transport and once at the posterior pole, the mRNPs are remodeled and Bruno dissociates, resulting in translation of *oskar*.

Altogether, our analysis of FMR1 function exemplifies how the protein binding landscape on RNAs and granule material properties work in concert to fine-tune RNA-protein interactions and post-transcriptional RNA regulation in a cellular context.

## Materials and Methods

### Fly husbandry and fly stocks

All flies were maintained at 25°C. The FMR1-GFP line used for co-localization, CLIP and iCLIP experiments was from Sudhakaran *et al.*^16^. FMR1^Δ50^ (BDSC_6930), FMR1^Δ113^ (BDSC_67403), FMR1-RNAi (BDSC_35200) and control RNAi (BDSC_35573) lines were obtained from the Bloomington Drosophila Stock Center. CRISPR injection and screening services at GenetiVision Corporation (http://www.genetivision.com/index.html) and protocols from https://flycrispr.org/ were used for generation of CRISPR flies. For CRISPR FMR1-sfGFP knock-in line, flies were injected with donor pHD-sfGFP-ScarlessDsRed-FMR1 (containing 1 kb homology arms of FMR1 from each side of the cut site) and gRNA: 5’- CTTCGATGGCACGTCCTAAGCCGAG-3’. For CTD deletion, pHD-sfGFP-ScarlessDsRed-FMR1-KH (containing KH domains only and CTD deleted), and the following two guide RNAs (used to make the deletion) were used:

gRNA1: 5’ - CTTCGTGACGCGGCGCTCTGAGCG - 3’

gRNA2: 5’ - CTTCGATGGCACGTCCTAAGCCGAG - 3’

### *oskar*-specific transcript pulldown and proteomics

*oskar*-specific pulldown to identify proteins associated with *oskar*-RNP complex was performed as described in Wippich and Ephrussi, 2020^35^, using the same probes and conditions, and the samples were sent for mass spectrometry.

### Mass spectrometry (LC-MS/MS) for *oskar*-specific transcript pulldown

Sample preparation: Cysteines were reduced with dithiothreitol at 56°C for 30 minutes (10 mM in 50 mM HEPES, pH 8.5) and further alkylated with 2-chloroacetamide at room temperature, in the dark for another 30 minutes (20 mM in 50 mM HEPES, pH 8.5). Samples were processed using the SP3 protocol^55^ and on-bead digested with trypsin (sequencing grade, Promega), which was added in an enzyme to protein ratio 1:50 for overnight digestion at 37°C. Peptides were modified with Isobaric Label Reagent (ThermoFisher) TMT10plex^56^ according the manufacturer’s instructions. For sample clean up, an OASIS® HLB µElution Plate (Waters) was used.

LC-MS/MS: An UltiMate 3000 RSLC nano LC system (Dionex) fitted with a trapping cartridge (µ- Precolumn C18 PepMap 100, 5µm, 300 µm i.d. x 5 mm, 100 Å) and an analytical column (nanoEase™ M/Z HSS T3 column 75 µm x 250 mm C18, 1.8 µm, 100 Å, Waters). Trapping was carried out with a constant flow of 0.05% trifluoroacetic acid at 30 µL/min onto the trapping column for 6 minutes. Subsequently, peptides were eluted via the analytical column with a constant flow of solvent A (0.1% formic acid in water) at 0.3 µL/min with increasing percentage of solvent B (0.1% formic acid in acetonitrile). The outlet of the analytical column was coupled directly to a QExactive plus (Thermo) mass spectrometer using the Nanospray Flex™ ion source in positive ion mode.

P0025: The peptides were introduced into the QExactive plus via a Pico-Tip Emitter 360 µm OD x 20 µm ID; 10 µm tip (New Objectives) and an applied spray voltage of 2.3 kV. The capillary temperature was set at 320°C. Full mass scan was acquired with mass range 350-1,500 m/z in profile mode with resolution of 70,000. The filling time was set at maximum of 10 ms with a limitation of 3x106 ions. Data dependent acquisition (DDA) was performed with the resolution of the Orbitrap set to 35000, with a fill time of 120 ms and a limitation of 2x105 ions. A normalized collision energy of 32 was applied. The isolation window of the quadrupole was set to 1.0 m/z. Dynamic exclusion time of 30 s was used. The peptide match algorithm was set to ‘preferred’ and charge exclusion ‘unassigned’, charge states 1, 5 - 8 were excluded. MS2 data was acquired in profile mode.

P0116 and P0163: The peptides were introduced into the QExactive plus via a Pico-Tip Emitter 360 µm OD x 20 µm ID; 10 µm tip (New Objectives) and an applied spray voltage of 2.3 kV. The capillary temperature was set at 320°C. Full mass scan was acquired with mass range 375-1,200 m/z in profile mode with resolution of 70,000. The filling time was set at maximum of 250 ms with a limitation of 3x106 ions. Data dependent acquisition (DDA) was performed with the resolution of the Orbitrap set to 35,000, with a fill time of 120 ms and a limitation of 2x105 ions. A normalized collision energy of 32 was applied. The isolation window of the quadrupole was set to 1.0 m/z. Dynamic exclusion time of 30 s was used. The peptide match algorithm was set to ‘preferred’ and charge exclusion ‘unassigned’, charge states 1, 5 - 8 were excluded. MS2 data was acquired in profile mode.

### Data analysis for *oskar*-specific transcript pulldown LC-MS/MS

IsobarQuant^57^ and Mascot (v2.2.07) were used to process the acquired data, which was searched against a Uniprot *Drosophila melanogaster* proteome database containing common contaminants and reversed sequences. The following modifications were included into the search parameters: Carbamidomethyl (C) and TMT10 (K) (fixed modification), Acetyl (Protein N-term), Oxidation (M) and TMT10 (N-term) (variable modifications). For the full scan (MS1) a mass error tolerance of 10 ppm and for MS/MS (MS2) spectra of 0.02 Da was set. Further parameters were set: Trypsin as protease with an allowance of maximum two missed cleavages: a minimum peptide length of seven amino acids; at least two unique peptides were required for a protein identification. The false discovery rate on peptide and protein level was set to 0.01.

### Single molecule fluorescence in-situ hybridization (smFISH)

Ovaries were dissected in PBS and fixed in 2% Paraformaldehyde in PBS + 0.1% Triton X-100 (PBT) for 20 min at RT on a rotator. They were washed twice with 750 µL PBT, 10 min each. 100 µL of HYBEC buffer (2x saline-sodium citrate (SSC), 15% ethylene carbonate, 1 mM EDTA, 50 µg/mL heparin, 100 µg/mL salmon sperm DNA, 1% Triton X-100) was then added to the samples and incubated at 42°C for 15 min. 100 µL of HYBEC buffer with 4 nM per probe concentration was then added to the 100 µL HYBEC with sample, and incubated for 2 hours at 42°C. The samples were washed as follows at RT: 10 min with HYBEC buffer at 42°C, 10 min with HYBEC:PBT (1:1) at 42°C, 10 min with PBT at 42°C and 10 min with PBT at RT. 100 µL of 80% 2,2’-thiodiethanol (TDE) in PBS was used as mounting medium. The samples were visualized using a Leica SP8 confocal microscope. Object based co-localization analysis was performed using the published R plugin – xsColoc as in Gaspar *et al.*^58^.

### UV crosslinking and immunoprecipitation (CLIP) for ovaries

Ovaries were harvested in PBS using a Kitchen Aid and resuspended in 600 µL lysis buffer (10 mM Tris-Cl (pH 7.5), 150 mM NaCl, 0.5 mM EDTA, 0.5% SDS, 0.5% NP-40, freshly added Roche protease inhibitor 1:100 and Ribolock 1:2000). The sample was split into two: one was used as a non-crosslinked control and other was crosslinked with 0.3 J/cm^2^ UV (254 nm) in a Stratalinker. Both samples were homogenized using a pestle and centrifuged for 5 min at 800 rpm, 4°C. The supernatant was diluted 1:1 (330 µL + 300 µL) with buffer A (10 mM Tris-Cl pH 7.5, 150 mM NaCl, 0.5 mM EDTA, freshly added Ribolock 1:2000 and Roche protease inhibitor 1:100). 5 µL GFP Trap magnetic agarose beads were added, and incubated for 2- 3 hours on rotator at 4°C. Beads were washed twice with wash buffer (10 mM Tris-Cl pH 7.5, 150 mM NaCl, 0.5 mM EDTA, 0.5% NP-40, 0.5% SDS, 0.02 mg/mL Heparin, freshly added Ribolock 1:2000 and Protease Inhibitor 1:100), twice with high salt wash buffer (10 mM Tris-Cl pH 7.5, 750 mM NaCl, 0.5 mM EDTA, 0.5% NP-40, 0.5% SDS, 0.02 mg/mL Heparin, freshly added Ribolock 1:2000 and Protease Inhibitor 1:100 and) and twice with wash buffer again, 10 min each on rotator at 4°C. The beads were resuspended in 100 µL Proteinase K Buffer (20 mM HEPES (pH 7.5), 150 mM NaCl, 1% SDS) and 0.2 mg/mL Proteinase K at 55°C for 45 min. The RNA was extracted using Trizol LS reagent, following the manufacturer’s instructions. First strand synthesis was performed using SuperScript™ III First-Strand Synthesis SuperMix for qRT-PCR following the manufacturer’s instructions, and the cDNA prepared was used for qRT-PCR using SYBR green qPCR mix in Step One Real Time PCR system from Applied Biosystems.

### Plasmids and cloning

Cloning of expression vectors for protein purification: cDNA synthesis was performed on RNA extracted from ovaries using SuperScript™ III First-Strand Synthesis SuperMix for qRT-PCR following the manufacturer’s instructions. FMR1 cDNA was amplified using the forward primer (FP) 5’- CACCACTACGTCTGGCGATATGGAAG-3’ and reverse primer (RP) 5’-GGACGTGCCATTGACCAG-3’ and cloned into pAW vector (pA-FMR1) using the Gateway Cloning System (https://emb.carnegiescience.edu/drosophila-gateway-vector-collection). To clone ΔN-FMR1 for *in vitro* competition assay, FMR1 was amplified from pA-FMR1 using FP 5’- AGACGGATCCATGGGAAACTACGTTGAGGAG-3’ and RP 5’-TCTGAGCTCTTAGGACGTGCCATTGACCAGGCC-3’. A pET11 vector and the amplicon were both digested with BamHI and SacI at 37°C for 30 min. The digested products were gel purified and ligated using T4 DNA ligase overnight at 16°C. The ligation mix was transformed into *E.coli* and positives were screened by sequencing. To clone ΔN-FMR1-GFP for *in vitro* iCLIP, 5’- AGCTTTCACTTGTAGAGCTCTTACTTGTAGAGCTCGTCCATG-3’ and 5’- CTGGTCAATGGCACGTCCGGCGGTATGGTGAGCAAGGGCGAGGA -3’ were used to amplify GFP from pA-EB1-GFP plasmid^59^. The amplicon was used as a primer with pET11-ΔN-FMR1 as vector to amplify and generate pET11-ΔN-FMR1-GFP. For cloning λN-sfGFP tagged FMR1 constructs, plasmid pMJ-His-TEV-lambdaN-sfGFP (gift from Mandy Jeske (Heidelberg University)) was digested with ScaI. Takara In-Fusion HD cloning kit was used as per manufacturer’s instructions, to generate different λN-tagged proteins. For λN-FMR1-KH-CTD, primers 5’-GTTCTGTTCCAGGGGCCCAGTATGGGAAACTACGTTGAGG-3’ and 5’- TCCGGTACCTCATTAAGTTTAAGTTTAGGACGTGCCATTGACCA-3’ were used; for λN-FMR1- KH, primers 5’-GTTCTGTTCCAGGGGCCCAGTATGGGAAACTACGTTGAGG-3’ and 5’- GTACCTCATTAAGTTTAAGTCTGATCAATCTCCATCTTCT-3’ were used and for λN-FMR1- CTD, primers 5’-CAACAGACCGGTGGATCCATGCAGCTTCGCGCCATCCAGGAA-3’ and 5’- TTCCTGGATGGCGCGAAGCTGCATGGATCCACCGGTCTGTTG-3’ were used. The DNA plasmid pFL-5xBoxB containing luc_BoxB reporter under T3 promoter was a gift from Mandy Jeske (Heidelberg University).

### Recombinant protein purification from *E. coli*

Electrocompetent *E. coli* bacteria were transformed with the expression vector and cultured at 37°C in LB media with appropriate antibiotics overnight. A fresh 1 L culture was started from the overnight inoculum and incubated at 37°C. When OD reached 0.5, 0.2 mM IPTG was used to induce protein expression. The culture was incubated overnight at 18°C. Cells were harvested by centrifugation for 20 min at 3000 rcf, 4°C. The pellet was resuspended in 20 mL water and centrifuged for 15 min at 3000 rcf, 4°C. The supernatant was removed and 20 mL of lysis buffer (500 mM NaCl, 20 mM Tris-Cl (pH 8.5), 40 mM imidazole, 5% glycerol, freshly added 5 mM beta-mercaptoethanol, 0.01% NP-40, 1X Roche protease inhibitor cocktail) was added to resuspend the pellet. A microfluidizer was used to lyse the cells. The sample was centrifuged for 20 min at 15000 rcf, 4°C in a Beckman SS34 rotor. The supernatant was collected. For His-tagged proteins, the lysate was passed through a His-Trap HP column. Sample was eluted in 2 mL volumes, using a 0–100% gradient of imidazole (600 mM) over a volume of 40 mL using elution buffer (500 mM NaCl, 20 mM Tris-Cl (pH 7.5), 5% glycerol, 600 mM imidazole). To cleave off the tag, the samples were incubated with protease overnight at 4°C. The protein sample was concentrated to 5 mL using Amicon Ultra centrifugal filters and injected onto a gel filtration column HiLoad 16/600 Superdex 200 pg. 1 mL fractions were collected using gel filtration buffer (150 mM NaCl, 20 mM Tris-Cl (pH 7.5), 2 mM MgCl_2_, 5% glycerol). The samples were then combined and concentrated.

### Western blot for ovaries

Four pairs of ovaries were dissected in PBS. 1X LDS sample buffer with 10 mM DTT (80 µl) was added and ovaries were crushed using a pestle. Samples were boiled for 10 min at 95°C. 20 µL of supernatant was loaded onto 4-12% bis-tris precast gel and run at 180 V for 1 hour. The gel was transferred to a nitrocellulose membrane with a semi-dry blotting apparatus from Bio-Rad. The membrane was blocked with 5% milk in TBST (Tris-buffered saline + 0.1% Tween 20) for 30 min at RT. The samples were incubated overnight in primary antibody at 4°C. The next day, the membrane was washed 10 min each (3x) in blocking buffer at RT, followed by incubation in secondary antibody for 2 hours at RT. The membrane was washed for 10 min each (3x) in TBST at RT. The membrane was developed using Immobilien Western HRP substrate peroxide solution (ECL). The following antibodies were used: mouse anti-FMRP (DSHB 5A11), rabbit anti-Osk (Ephrussi lab), rabbit anti-GFP (Torrey Pines Biolabs TP401), rabbit anti-Histone H3 (Abcam ab1791), mouse anti-Tubulin (Sigma Aldrich T607), Donkey ECL Anti-Rabbit IgG HRP linked whole antibody (GE Healthcare NA934), Sheep ECL Anti-Mouse IgG HRP linked whole Antibody (GE Healthcare NA931).

### qRT-PCR for ovaries

Three pairs of ovaries were dissected in ice cold PBS and 100 µL of Trizol LS reagent was added. The ovaries were ground using a pestle and 700 µL Trizol LS was added. 213 µL chloroform was added, vortexed and incubated on ice for 10 min. The sample was centrifuged at 13,200 rpm for 20 min at 4°C. The aqueous phase was extracted into a fresh tube and 0.5 µL of 15 mg/mL GlycoBlue^TM^, and 533 µL isopropanol per 750 µL Trizol LS were added and incubated for 10 min at RT. Samples were centrifuged at 13,200 rpm for 20 min at 4°C. The supernatant was removed. 1 mL 70% ethanol was added to the pellet and centrifuged at 13,200 rpm for 5 min at 4°C. The pellet was then air-dried and resuspended in 30 µL RNase/DNase free water. 3 µL of 10x DNase buffer and 1 µL Turbo DNase was added to the samples and incubated at 37°C for 30 min for DNA removal. 500 µL Trizol LS was then added, and the RNA extraction steps repeated. 1.5 µg of RNA was used for cDNA synthesis. Superscript III First-Strand Synthesis Supermix kit was used following manufacturer’s instructions. 2 µL of cDNA was then used for qPCR using SYBR Green PCR mix. Step One Real Time PCR system from Applied Biosystems was used, with the following conditions:

Step 1- 95°C, 10 min;

Step 2- 95°C, 15 sec;

Step 3- 60°C, 1 min;

(steps 2 and 3) cycle 40x.

*18S* RNA was used for normalization, and Ct values were calculated.

### Pole cell analysis

Flies were fed for 48 hours on yeast at 25°C and then transferred to cages with agarY plates (agar plus yeast) overnight. The next day, the plates were changed, and the flies were allowed to pre-lay for 1 hour on fresh agarY plates, after which the plates were discarded. The flies were put on fresh agarY plates for 2 hours, followed by changing the plates and allowing the plates with laid eggs to incubate for further 2 hours at 25°C. Embryos from were then collected from the plates and dechorionated using 50% bleach (sodium hypochlorite 6-14%) in water for 2 min. Embryos were washed with water. 500 µL of preheated (92°C) salt solution with 0.4% NaCl, 0.3% Triton X- 100 was added, and the embryos were heat fixed at 92°C for 30 sec. 1 volume ice cold salt solution was used to wash once, and 1 volume heptane plus 1 volume methanol (500 µL plus 500 µL) were added to the fixed embryos. Samples were vortexed for 30 seconds and embryos were allowed to sink for 10 seconds. Supernatant with floating embryos was discarded. The embryos were rinsed thrice with PBST (PBS + 0.1% Tween 20) and then washed thrice for 15 min each at RT in PBST. Embryos were blocked in blocking buffer (PBS, 0.3% Triton X-100, 0.5% BSA) for 1 hour at RT. They were then incubated in primary antibody (anti-Vasa rat antibody, Ephrussi lab) at 4°C overnight. The next day, the embryos were washed for 20 min twice in blocking buffer. They were blocked in blocking buffer B (PBS, 0.1% Triton X-100, 10% Normal Goat Serum) for 1 hour at RT, followed by secondary antibody (anti-rat Alexa Fluor 647) for 2 hours at RT. The secondary antibody was removed, and the embryos were stained with DAPI (1:2500 in PBS) for 5 min. Samples were washed for 20 min each twice with PBT (PBS, 0.1% Triton X-100) each. Embryos were mounted with Vectashield and visualized under a Leica confocal SP8 at 20x and the pole cells were counted.

### *In vitro* translation assay

Master mix (MM)-T (0.25 g/L tRNA, 0.05 M potassium acetate, 0.016 M HEPES-KOH, pH 7.5, 0.08 g/L creatine kinase, 2 nM reporter RNAs) was prepared for each reporter RNA (2 µL per reaction) and 2.5 µL of protein (gel filtration buffer for no protein control) was added to the reaction along with 40% *Drosophila* embryo extract (4 µL). The samples were pre-incubated for 20 min at 22°C. 1.5 µL ARS (0.1 mM amino acid mix, 0.02 M creatine phosphate, 0.8 mM ATP) was added to the reactions and samples were incubated for another 60 min at 22°C. The samples were then snap-frozen in liquid nitrogen to stop the reaction. A Mithras LB 940 plate reader was used to measure the luciferase activity as a readout for the protein levels in each sample. 50 µL of luciferase substrate (Promega, E1500) was dispensed into 4 µL of the reaction, it was shaken for 3 sec (2 mm orbital), and the reading was taken for 7 sec. The values were normalized to the no-protein control.

### In vivo iCLIP

Ovary extract preparation from FMR1-GFP and Flag-Myc-GFP (Ephrussi lab) fly lines: Newly hatched flies were transferred to a fresh vial with yeast and fed for 2 days at room temperature. The flies were collected and ground using the Kitchen Aid in ice cold PBS. The samples were sieved using 400 nm and 200 nm sieves, and the ovaries were collected using the 80 nm sieve. The ovaries were transferred to a 15 mL tube and centrifuged at 600 rcf for 30 sec at 4°C. The supernatant was removed and 12 mL PBS was added and samples centrifuged at 600 rcf for 30 sec at 4°C. The supernatant was removed and the ovaries were resuspended in 3 mL ice-cold PBS (one can get approximately 1.5 mL ovaries from 7.5 mL dry volume of flies). The ovaries were transferred to a 10 cm cell culture dish on ice. The ovaries were crosslinked with 254 nm UV using Stratalinker. The ovaries were collected in 15 mL tube, washed with PBS and centrifuged at 600 rcf for 30 sec at 4°C. The supernatant was removed and the ovaries were resuspended in 3 mL PBS and split into two 1.5 mL protein low binding tubes. The ovaries were centrifuged at 600 rcf for 30 sec at 4°C. The supernatant was removed, 200-300 µL iCLIP lysis buffer supplemented with DTT (1 mM) and Ribolock (1:1000 v/v) was added and the ovaries were lysed using a pestle. The lysate was centrifuged at 16,000 rcf for 10 min at 4°C. The supernatant was recovered and the centrifugation step was repeated. The supernatant was snap-frozen for storage at -80°C.

Bead preparation: For each sample, 10 µL of beads were used: for FMG CL α-GFP and FMR1 CL α-GFP we used Magnetic agarose GFP Trap beads (Chromotek); for FMR1 CL beads-only Magnetic agarose beads (Chromotek) were used. The beads were washed 2x with 1 mL iCLIP high salt buffer (50 mM Tris-HCl, pH 7.4, 1 M NaCl, 1 mM EDTA pH 8.0, 1% Igepal CA-630, 0.1% SDS, 0.5% sodium deoxycholate, freshly added 1 mM DTT and 0.025 mg/mL heparin), and resuspended in 1 mL iCLIP lysis buffer (50 mM Tris-HCl pH 7.4, 100 mM NaCl, 1% Igepal CA- 630, 0.1% SDS, 0.5% Sodium deoxycholate, supplemented with 1mM DTT, Ribolock (1:2000 v/v)). For each sample in the experiment, 1,030 µL sample aliquot with 20 mg of total protein content was used. iCLIP lysis buffer supplemented with 1mM DTT and Ribolock (1:1000 v/v) was used for diluting the extracts. 5 µL Turbo DNase was added with 10 µL RNase I (Invitrogen; diluted 1:20) and incubated at 37°C for 3 min at 1,100 rpm. The samples were incubated on ice for >3 min. 10 µL of the prepared beads were then added to each sample.

Immunoprecipitation: The samples were incubated for 2 h in cold room on a rotating wheel (11 rpm). The samples were washed 3x with 1 mL iCLIP high salt buffer, transferred to new tubes, and then washed 3x with PNK wash Buffer BA. The RNA 3’end dephosphorylation and the subsequent steps until sequencing were performed as mentioned in the "*in vitro* iCLIP" method. 10 different second adapters (L01clip2.0 to L10clip2.0) were used for library preparation as follows:

| Replicate | Sample | Primer |  |
| --- | --- | --- | --- |
| 1 | FMG X-linked anti-GFP | L01clip2.0 | CGTGAT |
| 1 | FMR1 X-Linked Beads only | L02clip2.0 | ACATCG |
| 1 | FMR1 X-Linked anti-GFP | L03clip2.0 | GCCTAA |
| 2 | FMG X-linked anti-GFP | L04clip2.0 | TGGTCA |
| 2 | FMR1 X-Linked Beads only | L05clip2.0 | CACTGT |
| 2 | FMR1 X-Linked anti-GFP | L06clip2.0 | ATTGGC |
| 3 | FMG X-linked anti-GFP | L07clip2.0 | GATCTG |
| 3 | FMR1 X-Linked Beads only | L08clip2.0 | TCAAGT |
| 3 | FMR1 X-Linked anti-GFP | L09clip2.0 | CTGATC |
| 4 | FMG X-linked anti-GFP | L10clip2.0 | AAGCTA |
| 4 | FMR1 X-Linked Beads only | L01clip2.0 | CGTGAT |
| 4 | FMR1 X-Linked anti-GFP | L02clip2.0 | ACATCG |

The *in vivo* iCLIP libraries were sequenced on an Illumina HiSeq 2500 machine as 62 nt single-end reads including a 6 nt sample barcode as well as 5+4 nt unique molecular identifiers (UMIs).

### In vitro iCLIP

Sample preparation and immunoprecipitation: *In vitro* transcription (IVT) of *oskar* mRNA and *bcd* 3’UTR (to be used as spike-in) was performed using a MEGAscript T7 transcription kit (Invitrogen) according to the manufacturer’s instructions. Briefly, template for IVT was prepared by PCR using T7-forward primer 5’-GGATCACTTTCCTCCAAGCG-3’ and *oskar*-specific reverse primer 5’- CCTATAACAAGCTGCAATGTAAAATCC-3’. 200 ng template DNA was used for a 20 μl transcription reaction for 3 h at 37°C. Template DNA was digested with Turbo DNase. The RNA was extracted using phenol chloroform as follows - 15 μl of Ammonium-o-acetate stop solution and 115 μl of nuclease free water was added to the samples and mixed well. Equal volume of phenol/chloroform (acidic, Ambion) was added, vortexed thoroughly, followed by equal volume of chloroform. The samples were centrifuged at 12,000 rcf for 15 min at 4°C. The aqueous phase was extracted, and RNA was precipitated using two volumes of 100% ice cold ethanol. The samples were centrifuged at 12,000 rcf for 30 min at 4°C. The supernatant was discarded and the pellets were washed with 1 mL of 70% ethanol followed by centrifugation at 12,000 rcf for 5 min at 4°C. The pellets were then air dried for 5 min and resuspended in 20 μL nuclease-free water.

*In vitro* transcribed *oskar* RNA and *bcd* 3’UTR were heated to 70°C for 5 min and immediately put on ice. 100 nM or 250 nM purified recombinant protein ΔN-FMR1 was incubated with 20 nM RNA in binding buffer BA for 10 min at 37°C at 1100 rpm. The samples were put on a sterile 10 x 10 cm plate and crosslinked with 254 nm UV at 6 mJ/cm^2^ in a Stratalinker. The volume was made up to 900 µL with lysis buffer BA, and 1:650 diluted RNase I and 1 µL Turbo DNase were added to the samples. Samples were incubated at 37°C for 3 min, followed by addition of 15 µL GFP Trap magnetic beads (Chromotek) and incubation at 4°C for 1 hour on a rotator (15 rpm). The beads were washed 3x with high-salt wash buffer BA, and thrice with PNK wash buffer BA, 10 min each on rotator (20 rpm).

RNA 3’end dephosphorylation: The beads were resuspended in 20 µL of PNK reaction mix and incubated at 37°C for 20 min at 1100 rpm. They were washed 1x with PNK wash buffer, 2x with high salt wash buffer BA, and 2x with PNK wash buffer BA, 10 min each on rotator.

First adapter ligation to the 3’end of RNA: The beads were resuspended in 20 µL adapter ligation reaction mix and incubated overnight at 16°C at 1100 rpm. 500 µL PNK wash buffer was added and the beads washed 2x with high salt wash buffer BA, and 2x with PNK wash buffer BA (tubes were changed after first PNK wash buffer wash) for 10 min each on rotator.

Radioactive labelling of RNA 5’end: To radioactively label the bound RNAs, the beads were resuspended in 16 µL of Hot PNK mix and incubated at 37°C for 5 min at 1,100 rpm. The beads were washed 1x with PNK wash buffer BA, followed by addition of 20 µL 1X LDS sample buffer, and incubation at 70°C for 5 min. The supernatant was run on a 4-12% Bis-Tris precast gel in MOPS running buffer. The gel was exposed to a phosphor screen (Fujifilm), and the screen was visualized using a Typhoon FLA 9500 biomolecular imager from GE healthcare. Using the autoradiograph as mask, the membrane was cut and put into tubes to extract the RNA. 10 µL of proteinase K (20 mg/mL) in 200 μL proteinase K (PK) buffer was added to the membrane and incubated at 37°C for 20 min at 1,100 rpm. 200 µL of PK + urea buffer was then added and incubated for 20 min at 37°C, 1100 rpm. The solution was put in a Phase Lock Gel Heavy tube along with 400 μL of phenol/chloroform (pH 7.8/8.0). The samples were incubated at 30°C, 1100 rpm for 5 min, and the phases were separated by centrifuging at 16,000 rcf for 5 min at RT. The aqueous layer was transferred to a new tube, and the RNA was precipitated by the addition of 0.75 µL GlycoBlue (15 mg/mL), 40 μL 3 M sodium acetate (pH 5.5), and 1 mL 100% ethanol. The solution was mixed by inverting the tube and incubated at -20°C overnight. The next day, the samples were centrifuged at 21,100 rcf for 20 min at 4°C, washed 1x with 80% ethanol, centrifuged for 5 min and the pellets air dried for 3 min. Pellets were resuspended in 5 μL ultrapure water.

Reverse transcription: 1 µL of RT-oligo (0.5 pmol/μL) and 1 µL of dNTP mix (10 mM) were added to the samples and incubated at 70°C for 5 min, followed by the 13 µL of RT-CLP mix. The following RT program was run: 25°C 5 min, 42°C 20 min, 50°C 40 min, 80°C 5 min, 4°C hold. 1.65 μL of 1 M NaOH was added and incubated at 98°C for 20 min, followed by addition of 20 µL 1 M HEPES-NaOH (pH 7.3).

MyONE Silane cleanup: 10 µL MyONE silane beads per sample were washed and resuspended in 93 µL RLT buffer and added to the samples. 112 μL 100% ethanol was added and mixed by pipetting, and incubated for 5 min at RT. The beads were mixed again and incubated for 5 min at RT. The beads were magnetically separated, and the supernatant was discarded. The beads were resuspended in 1 mL of 80% ethanol and transferred to a new tube. The step was repeated 2x, and the beads were air-dried for 5 min at RT. The beads were resuspended in 5 µL ultrapure water, and incubated for 5 min at RT.

Second adapter ligation: To the cDNA-bead solution, 2 µL of L##clip2.0 (second adapter) and 1 μL 100% DMSO were added, incubated at 75°C for 2 min and then immediately put on ice. Since there were a total of 7 samples, 7 different second adapters (L07clip 2.0, L08clip 2.0, L09clip 2.0 and L10clip 2.0 for 100 nM ΔN-FMR1; L12clip2.0, L13clip2.0 and L14clip2.0 for 250 nM ΔN-FMR1) were used. 12 µL of Lig-CLP mix was added to each sample and mixed well, followed by the addition of 1 µL RNA ligase. The samples were then incubated at RT, 1100 rpm overnight. MyONE Silane cleanup: The next day, 5 µL silane beads per sample were washed and resuspended in 60 µL RLT buffer. The 60 µL beads were added to each sample along with 60 µL 100% ethanol, mixed by pipetting and incubated at RT for 5min. The step of mixing and incubation was repeated and the beads magnetically separated. The beads were then resuspended in 1 mL 80% ethanol and transferred to a new tube. The step was repeated 2x, and the beads were air-dried for 5 min at RT, resuspended in 23 µL ultrapure water and incubated for 5 min at RT.

First PCR amplification: 2X Phusion high fidelity master mix (25 µL) and P5Solexa_S and P3Solexa_S primer mix (2.5 µL of 10μM stock) were mixed and added to 22.5 µL cDNA sample. The following PCR program was used: 98°C 30 sec 1x, 98°C 10 sec 6x, 65°C 30 sec, 72°C 30 sec, 72°C 180 sec, 16°C hold

First ProNex Size selection: ProNex chemistry was equilibrated to RT for 30 min and the beads were vortexed. 1:2.95 v/v ratio of sample:beads was added to the samples. They were mixed by pipetting up and down 10 times and incubated at RT for 10 min. The samples were placed on a magnetic rack and let stand for 2 min. The supernatant was removed and 500 µL of ProNex wash buffer was added to the beads still on the rack. The samples were incubated for 30-60 sec and the buffer was removed. The step was repeated once, and the samples were air-dried for 8-10 min until the beads began to crack. The samples were removed from the rack, and 23 µL water was added to elute the samples. The samples were incubated at RT for 5 min. The beads were then put back on the magnetic rack and the supernatants carefully transferred to a new tube.

Test PCR amplification: 9 µL of PCR mix was prepared- 2X Phusion high fidelity master mix (5 µL) and P5Solexa_S and P3Solexa_S primer mix (0.5 µL of 10 μM stock), and 3.5 µL water; and added to 1 µL cDNA. The following PCR program was run: 98°C 30 sec 1x, 98°C 10 sec 6x and 9x, 65°C 30 sec, 72°C 30 sec, 72°C 180 sec, 16°C hold. 2 µL of the amplified libraries were run on capillary gel electrophoresis using the High Sensitivity D1000 kit in a TapeStation system, to test if the libraries look good.

Preparative PCR: The same conditions of test PCR amplification were repeated for 10 µL cDNA. 2 µL was again analyzed using capillary gel electrophoresis, and if all looked well, the second half of the libraries were also amplified.

Second ProNex size selection: The ProNex size selection step was repeated with 1:2.4 v/v ratio of sample:beads, and the samples were eluted in 72 µL of water. The concentration of the libraries was calculated using Qubit hsDNA kit, and were diluted to 10 nM, 20 µL for sequencing. The libraries were sequenced on an Illumina NextSeq 500 machine as 166 nt single-end reads including a 6 nt sample barcode as well as 5+4 nt unique molecular identifiers (UMIs).

|  |  |
| --- | --- |
| Lysis Buffer BA | 50 mM Tris-HCl, pH 7.4, 100 mM NaCl, 1% Igepal CA-630, 0.1% SDS, 0.5% sodium deoxycholate |
| Binding Buffer BA | 100 mM HEPES (pH 7.2), 30 mM MgCl<sub>2</sub>, 30% glycerol, 10 mM DTT, 100 mM KCl |
| High-salt wash Buffer BA | 50 mM Tris-HCl, pH 7.4, 1 M NaCl, 1 mM EDTA pH 8.0, 1% Igepal CA-630, 0.1% SDS, 0.5% sodium deoxycholate, freshly added 1 mM DTT and 0.025 mg/mL heparin |
| PNK wash Buffer BA | 20 mM Tris-HCl, pH 7.4, 10 mM MgCl<sub>2</sub>, 0.2% Tween 20, freshly added 1 mM DTT |
| 5X PNK Buffer, pH6.5 | 350 mM Tris-HCl, pH 6.5, 50 mM MgCl<sub>2</sub>, 5 mM DTT |
| PNK reaction mix | 15 µL water, 4 µL 5X PNK buffer (pH 6.5), 0.5 µL Ribolock, 0.5 µL T4 PNK enzyme |
| Adapter ligation reaction mix | 8 µL water, 5 µL 4x ligation buffer, 0.5 µL Ribolock, 1.5 µL preadenylated L3-App (20 µM), 4 µL PEG400, 1 µL T4 RNA Ligase |
| 4X Ligation Buffer | 200 mM Tris-HCl, pH 7.8, 40 mM MgCl<sub>2</sub>, 4 mM DTT |
| Hot PNK mix | 6 µL water, 0.8 µL 10X PNK buffer, 0.8 µL <sup>32</sup>P-gATP, 0.4 µL T4 PNK (New England Biolabs) |
| Proteinase K (PK) Buffer B | 100 mM Tris-HCl, pH 7.4, 50 mM NaCl, 10 mM EDTA |
| PK + urea Buffer | 100 mM Tris-HCl, pH 7.4, 50 mM NaCl, 10 mM EDTA, 7 M urea |
| RT-CLP mix | 7 µL water, 4 µL 5X RT buffer, 1 µL 0.1 M DTT, 0.5 µL Ribolock, 0.5 µL Superscript III |
| Lig-CLP mix | 2 µL 10X RNA ligase buffer (New England Biolabs), 0.2 µL 0.1 M ATP, 9 µL 50% PEG8000, 0.3 µL water, 0.5 µL high conc. RNA ligase |

### *In vitro* competition assay

The assay was adapted from Zarnack *et al.*^60^. Oligonucleotide (BRE A’_II: 5’- UUUAUUUAUAUGUUCGUGCACUUGUCCUAG-3’) was radioactively labeled with γ-^32^P-ATP using PNK as follows: 1 µL of oligo (10 μM) was added to 39 µL of UV buffer (1x PNK buffer A (Thermo Scientific), 10 μM DTT, 300 μCi γ-^32^P-ATP, 30 units PNK enzyme (Thermo Scientific)). The reaction was incubated for 1 h at 37°C, followed by 95°C for 2 min. 9 µL of unlabeled 10 μM oligo was added with water to make up the volume to 100 µL. To remove unincorporated γ-^32^P- ATP, the sample was passed through G-25 filter columns (GE Healthcare). 100 nM of labelled probe was incubated with 2 µM ΔN-FMR1. Reaction samples with recombinant Bruno protein (gift from Mainak Bose) in binding buffer (10 mM Tris (pH 7.4), 100 mM KCl, 2.5 mM MgCl_2_) were prepared in different concentrations and added to the oligo-ΔN-FMR1 mix such that final volume is 20 µL. The reactions were incubated for 15 min at 37°C, and then UV (254 nm) crosslinked at 150 mJ/cm^2^ using Stratalinker 2400. 4x SDS loading dye with 100 mM DTT was added to the samples and incubated to 95°C for 5 min. A NuPAGE Bis-Tris 4-12% precast gel was run, and the gel was first exposed to Fuji film phosphor screen and then stained with Coomassie. The phosphor screen was visualized using Typhoon FLA 9500 from GE healthcare.

### iCLIP data processing

Basic quality checks were done using FastQC (v0.11.8) (https://www.bioinformatics.babraham.ac.uk/projects/fastqc/) and reads were filtered based on sequencing qualities (Phred score) in the barcode and UMI regions using the FASTX-Toolkit (v0.0.14) (http://hannonlab.cshl.edu/fastx_toolkit/) and seqtk (v1.3) (https://github.com/lh3/seqtk/). Reads with a Phred score below 10 in the considered regions were removed from further analysis. Reads were de-multiplexed based on the sample barcode, which is found on positions 6 to 11, using Flexbar (v3.4.0) ^61^. Barcode and UMI regions as well as adapter sequences were trimmed from read ends using Flexbar requiring a minimal overlap of 1 nt of read and adapter and adding UMIs to the read names. Reads shorter than 15 nt were removed from further analysis. The downstream analysis was done as described in Chapters 3.4, 4.1 and 4.2 of ^62^. While only canonical chromosomes (2L, 2R, 3L, 3R, 4, X, Y, and mitochondrion_genome) of the *Drosophila melanogaster* genome assembly (BDGP6.28) and annotation of Ensembl ^63^ release 102 were used during mapping of *in vivo* iCLIP data, all chromosomes and scaffolds of the *Drosophila melanogaster* genome assembly (BDGP6.32) and annotation of Ensembl release 104 were used when mapping *in vitro* iCLIP data.

### Identification of binding sites

For *in vivo* iCLIP, BAM files from four FMR1 iCLIP replicates were merged and indexed using SAMtools (version 1.23) ^64^. The merged data were used as input for PureCLIP (version 1.0.3) ^37^ using *Drosophila melanogaster* reference genome BDGP6.28 (FASTA format). For binding site definition, the PureCLIP-called crosslink sites (BED format) together with strand-specific crosslink signal tracks (BigWig format) from the four FMR1 iCLIP replicates were used as input for the R/Bioconductor package BindingSiteFInder (version 2.8.0; https://bioconductor.org/packages/BindingSiteFinder/ ^62^ ^38^). Only standard chromosomes with the mitochondrial chromosome were retained. Gene and transcript annotations were extracted from *Drosophila melanogaster* BDGP6.28.102 (GTF format) using the R/Bioconductor packages txdbmaker (version 1.6.2;^65^, rtracklayer (version 1.70.1) ^66^, and GenomicFeatures (version 1.62.0) ^67^. Binding sites were identified using the function BSFind() with parameters est.subsetChromosome=“2L”, cutoff.geneWiseFilter=0.5, and match.geneType=“gene_biotype”. The optimal binding sites width determined by binding site finder was 7 nt. During reproducibility filtering, binding sites were kept if present in at least three out of four replicates with a 5% quantile cutoff (cutoff=0.05, nReps=3). Binding sites were assigned to gene biotypes in the following hierarchy: protein-coding > snRNA > snoRNA > ncRNA > pre-miRNA > tRNAs > rRNAs > pseudogene (overlaps.geneAssignment=“hierarchy”). In total, we identified 160,960 *in vivo* FMR1 binding sites on the transcripts of 3,677 genes.

For *in vitro* iCLIP on *oskar* mRNA, seven replicates were used for binding site identification (4x 100 nM and 3x 250 nM ΔN-FMR1-GFP). Initial peaks were defined on the merged crosslink signal track (BigWig format) as sites with crosslink signal exceeding ∼244, empirically determined based on the crosslink signal distribution. These were used together with the crosslink signal tracks from the replicates (BigWig format) as input for BindingSiteFinder. Binding sites were identified using BSFind() with a fixed binding site width of 7 nt (bsSize=7), as determined on the *in vivo* data, and parameters cutoff.geneWiseFilter=0.9 and overlaps.geneAssignment=“keep”. During reproducibility filtering, binding sites were kept if present in at least six out of seven replicates with a 5% quantile cutoff (cutoff=0.05, nReps=6). This procedure yielded a total of 41 *in vitro* FMR1 binding sites on *oskar* mRNA.

## Statistics and reproducibility

For all quantifications, statistical analyses were performed and data were plotted using Prism 11. P-value of <0.05 was considered significant. The statistical tests used and P-values are indicated in the figure legends. Sample sizes used were similar to those reported in previous publications. The number of replicates and sample sizes are indicated in the respective figure legends.

## Data Availability

The mass spectrometry proteomics data have been deposited at OSF, and the sequencing data for iCLIP have been deposited at the NCBI Gene Expression Omnibus (GEO). The links and accession numbers will be made available upon publication.

## Acknowledgements

We are grateful to Mandy Jeske (Heidelberg University) for help with the *in vitro* translation assay and for providing plasmids, to Mainak Bose for sharing recombinant Bruno protein, and to Alessandra Reversi for *Drosophila* transgenesis. We thank the EMBL Proteomics Core Facility for help with the *oskar* transcript-specific pulldown and associated MS. We thank Mainak Bose, Florence Besse and Imre Gaspar for discussions and careful reading of the manuscript. Stocks obtained from the Bloomington Drosophila Stock Center (NIH P40OD018537) were used in this study. M.B. was supported by an EMBL International PhD Program fellowship. This work was supported by DFG-FOR 2333 grants of the Deutsche Forschungsgemeinschaft (DFG) to A.E. (EP 37/2-1, 4-1), K.Z. (ZA 881/3-1) and J.K. (KO 4566/5-1). This work was also funded by the DFG via EXC 3113/1, Cluster for Nucleic Acid Sciences and Technologies – NUCLEATE (project ID 533767322), to K.Z and J.K. A.E. acknowledges funding from the EMBL. We acknowledge the IMB Genomics Core Facility and its NextSeq 500 sequencer (funded by the Deutsche Forschungsgemeinschaft [DFG, German Research Foundation] P#329045328).

## Author Contributions

V.G. and A.E. conceived the study, interpreted the results and wrote the manuscript. V.G. designed and performed the experiments and analyzed the data. F.W. performed the *oskar* transcript-specific pulldown and CLIP for FMR1. M.R. analyzed the mass spectrometry data. M.B. performed the *in vivo* iCLIP. V.G. and A.O. performed the *in vitro* iCLIP. D.L., A.B., A.L., J.K. and K.Z. analyzed the iCLIP data.

---

**Supplementary Figure S1:**
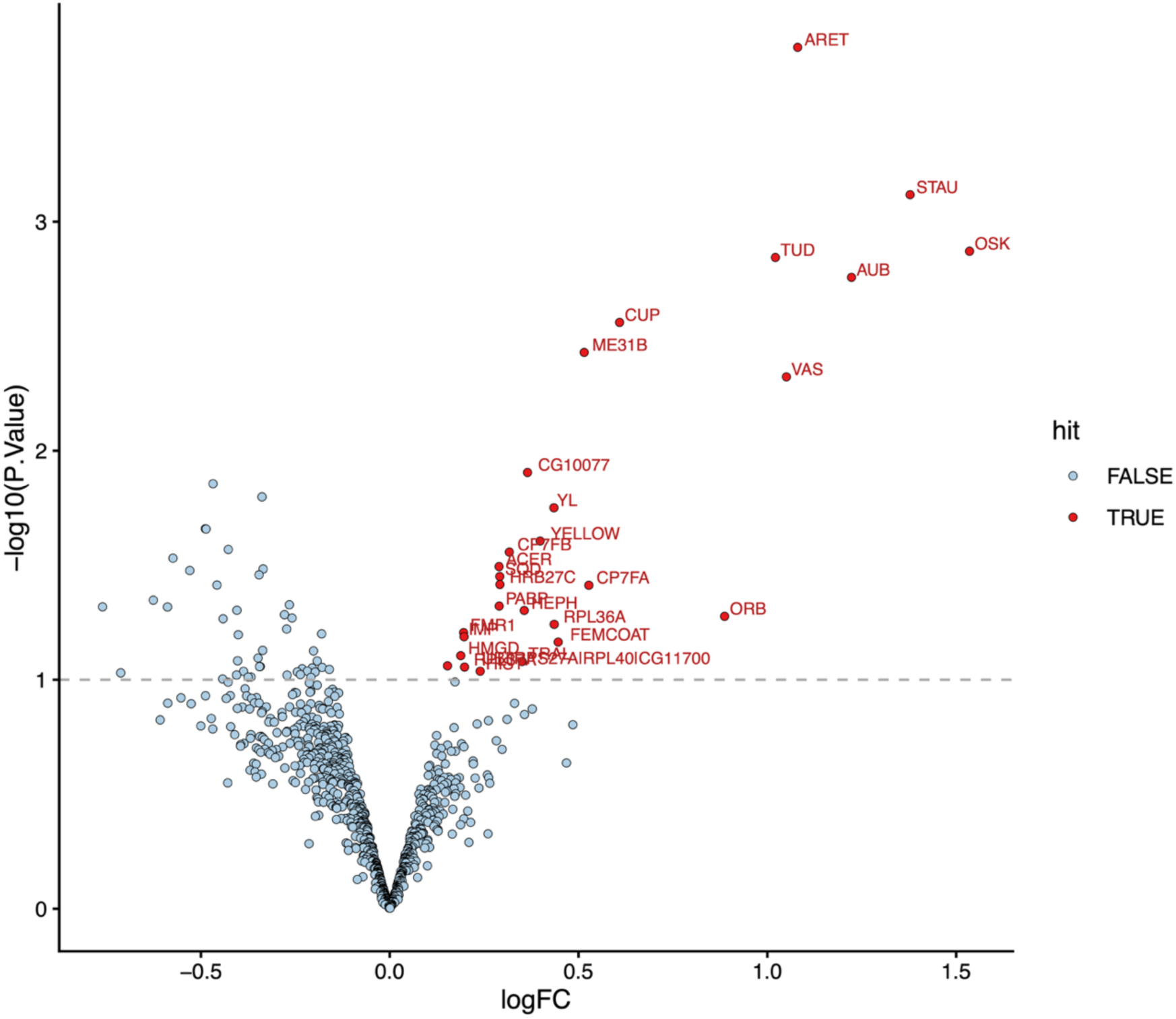
Proteins associated with *oskar* mRNA as identified in *oskar*-specific mRNP pulldown. Volcano plot showing proteins significantly enriched in *oskar*-specific transcript pulldown

**Supplementary Figure S2:**
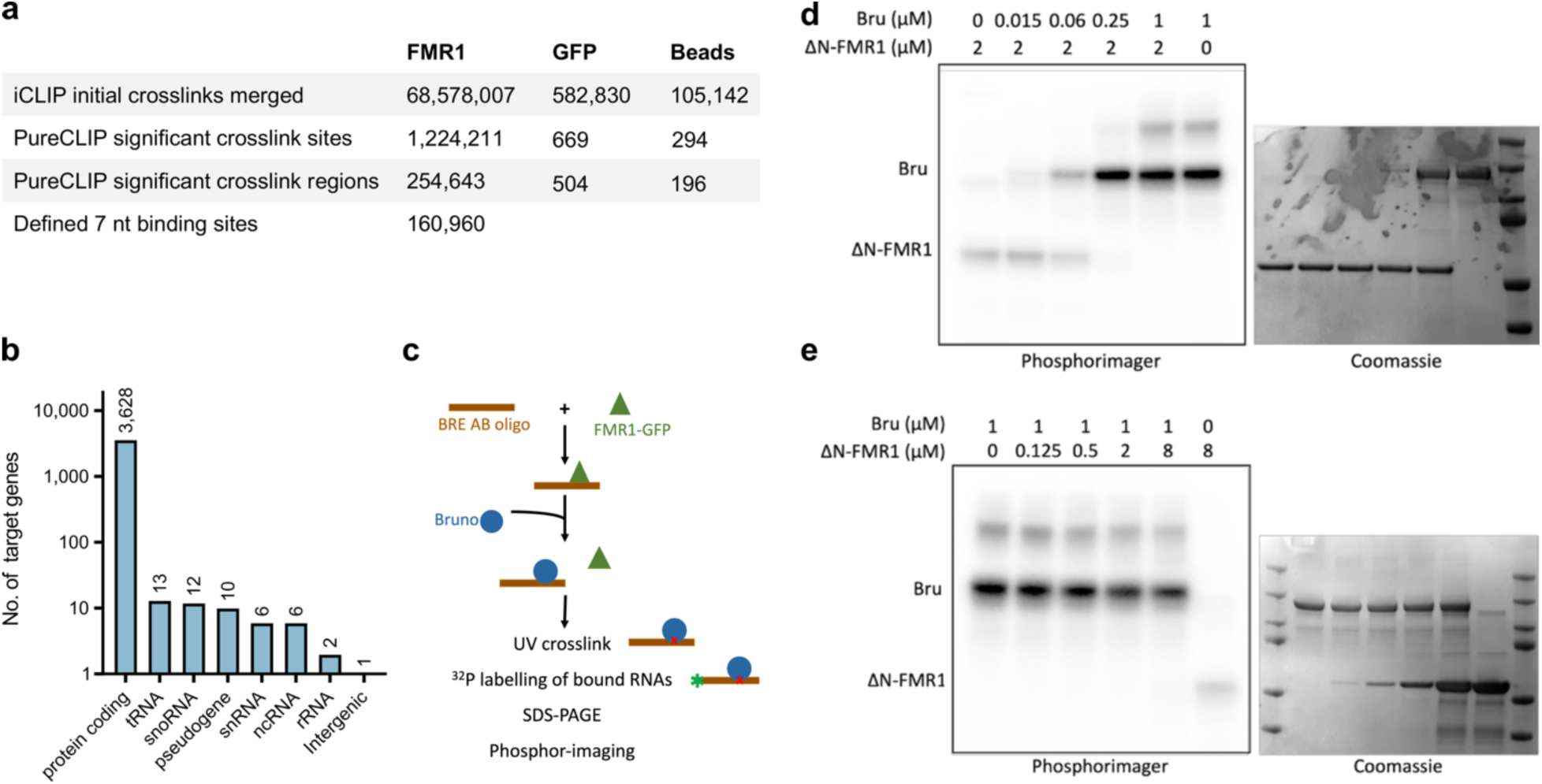
Bruno exhibits a higher affinity than FMR1 for BRE AB sites *in vitro*. **a**. Table shows the number of crosslink sites initially identified, after PureCLIP peak calling, and final binding sites identified in the three sample types used. **b**. Graph shows the classification of FMR1 target genes identified by *in vivo* iCLIP. **c**. Schematic for the competition assay between FMR1 and Bruno for BRE AB sites. **d**. Adding increasing concentrations of Bruno to ΔN-FMR1-BRE AB complex displaced FMR1 from the complex, indicating a higher affinity of Bruno for BRE AB. **e**. ΔN-FMR1, on the other hand, could not outcompete Bruno even at 8 times higher concentration. Coomassie staining is used for loading control of the two proteins.

**Supplementary Figure S3:**
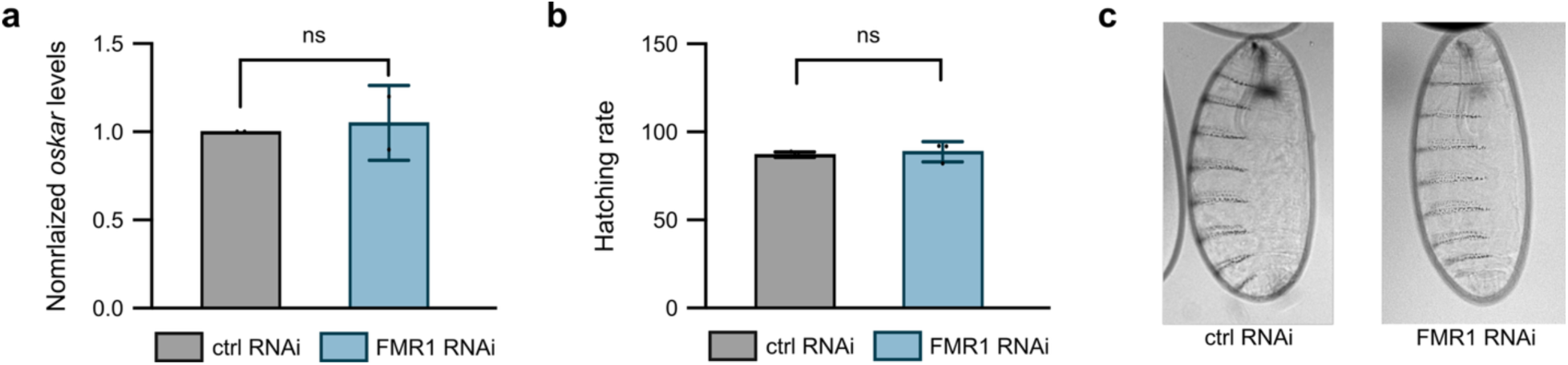
FMR1 knockdown has no effect of *oskar* mRNA levels, hatching rate or embryonic patterning. **a**. qRT-PCR analysis of *oskar* mRNA levels in control and FMR1 RNAi lines. *oskar* mRNA levels were normalized to *18S* RNA. **b**. Knockdown of FMR1 had no effect on the hatching rates of embryos, as compared to control. **c**. Knockdown of FMR1 had no effect on embryonic patterning of embryos, as compared to controls. Error bars represent standard deviation of the mean, n = 3 biological replicates. Student’s *t*-test was used for statistical analysis. P-values: ****<0.0001, ***<0.0005, **<0.005.

